# CD24 and CD27 resolve ontogeny and gradual differentiation of CD11c+ atypical memory B cells after malaria infection

**DOI:** 10.1101/2025.09.15.676235

**Authors:** Alan-Dine Courey-Ghaouzi, Linn Kleberg, Zaynab Mousavian, Maximilian Julius Lautenbach, Marta Pirronello, Ganesh E. Phad, Mattias N. E. Forsell, Anna Färnert, Christopher Sundling

## Abstract

Atypical B cells (ABCs) are observed across infections, yet their ontogeny and differentiation remain unclear. Using high-dimensional flow cytometry, single-cell transcriptomics, V(D)J sequencing, and functional assays in human malaria and healthy donors, we identified markers of cellular origin while tracking ABC differentiation. Early ABCs express CD11c and arise predominantly from memory B cells. Two intermediate states originate from germinal center-derived and marginal zone-like B cells. Pseudotime and lineage analyses reveal trajectories marked by progressive CD24 and CD27 loss, recapitulated in vitro, defining bona fide differentiation programs. Differentiation culminates in late ABCs lacking CD24, CD27, and CD21, with high CD11c, T-bet, and NKG7, partly overlapping with double-negative 2 cells. Early- and intermediate stages show greater plasma-cell differentiation potential and cytokine production capacity, whereas later stages acquire antigen-presentation transcriptomic signatures and microbial-particle association, defining stage-dependent functions that reconcile conflicting ABC roles. Comparable ABC populations occur in healthy blood and tonsils, with tissue-associated class switching and CD86 expression consistent with recent T-cell interaction. This framework consolidates ABC definitions onto a single differentiation axis and clarifies cellular origins.

**Graphical abstract:** 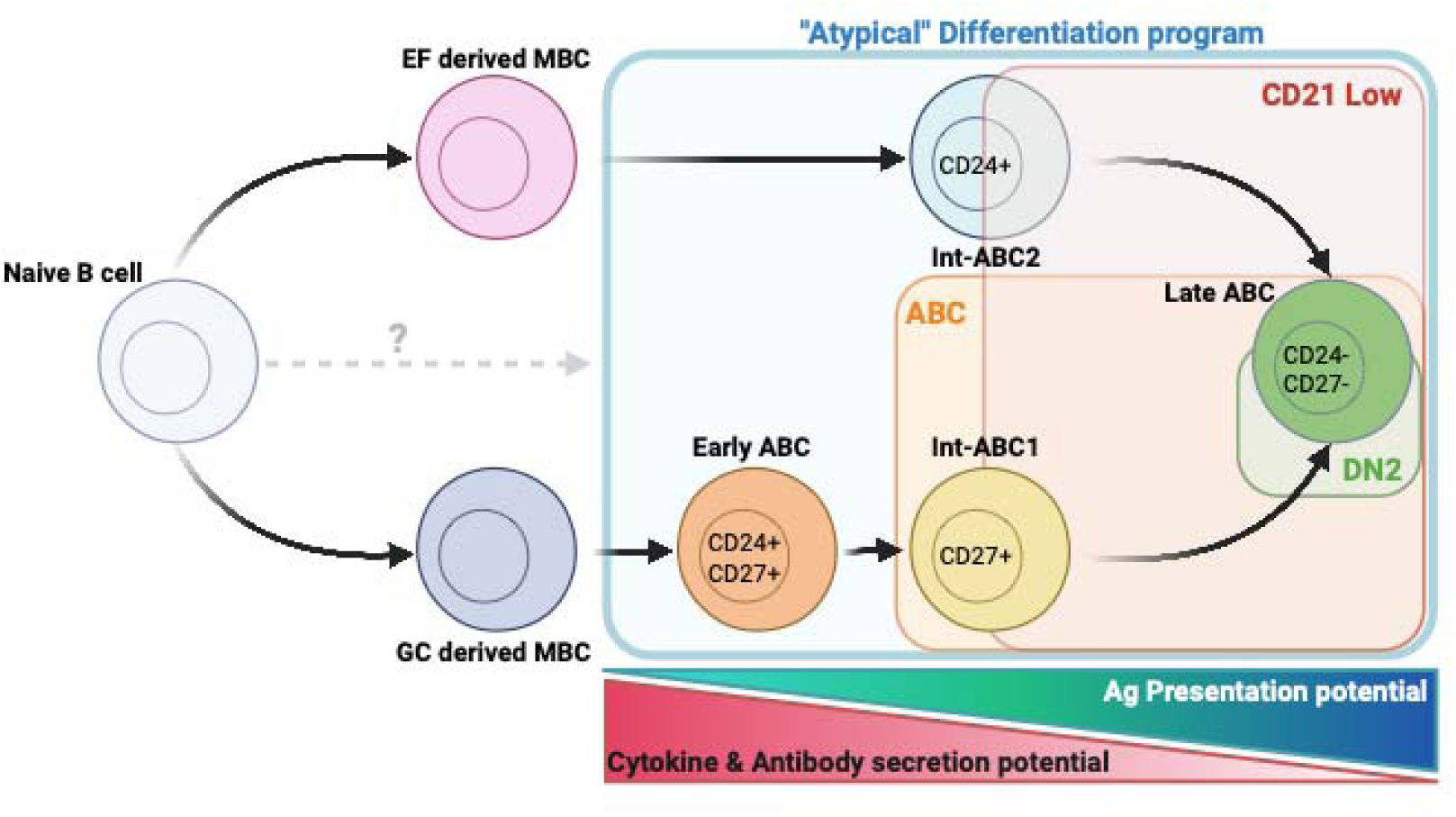

## Introduction

B cells are central players in adaptive immunity, performing essential roles such as antibody production, memory formation, and regulation of immune responses. Among the diverse subsets of B cells, atypical B cells (ABCs) have gained increasing attention due to their association with different disease states and have distinct functional and phenotypic profiles (*1–3*). Initially described as “exhausted” because of their altered phenotypes with elevated expression of immunomodulatory receptors and reduced BCR dependent responsiveness (*4, 5*), ABCs are now increasingly recognized as a dynamic population with context dependent marker expression and functions.

ABCs can be defined using several combinations of markers. A minimal definition uses loss of CD21 and CD27 (CD21– CD27–) together with CD11c expression (*6*). Many studies add T-bet (*7*), and sometimes FCRL5 (*8*) or CXCR3 (*9*), for greater specificity. Others use gating for IgD– CD27– double negative (DN) cells, with DN2 denoting the T-bet+ CD11c+ fraction (*10*). These definitions are not equivalent and identify overlapping but not identical populations: CD21– CD27– gates capture cells beyond T-bet+ CD11c+ ABCs, while DN2 or other multi marker definitions exclude some CD21– CD27– cells and IgD+ ABCs. Antigen specific approaches enrich class switched, high affinity cells and underrepresent IgM+ or IgD+ subsets. Marker thresholds, panel composition, and tissue context also vary, and terminology (atypical, age associated, T-bet+, DN2) is applied inconsistently. This limits comparability across studies, although efforts have been made to establish a standardized nomenclature (*11*). Additional variability arises from differential expression of markers such as FcRL5 and FcRL4 in malaria or HIV, respectively (*12, 13*). It remains unclear whether this variability stems from differences in gating strategies, reflects distinct stages of cellular differentiation, or perhaps site of differentiation.

Two recent field-wide reviews synthesize these issues and converge on differentiation as one of the central questions (*14, 15*). Taken together, they argue that ABCs are best understood as activated states that multiple B cell precursors can enter under distinct inflammatory milieus. A key area that remains unresolved is the relative contribution of naïve versus memory origins in humans, the anatomical settings from where the cells are generated (germinal center versus germinal center-independent/extrafollicular), and the signal combinations, including BCR engagement with IFNγ and TLR cues and differing modes of T cell help, which imprint CD11c/T-bet programs without uniquely specifying ontogeny. Equally unresolved are their temporal fates, i.e., whether these activated states are transient effectors, plasmablast precursors, or longer-lived memory across human timescales. Both reviews caution that common phenotypic gates (e.g., CD21low/CD27−; CD11c+/Tbet+) are insufficient to assign origin and recommend reporting, alongside phenotype, the inferred precursor, anatomical context, BCR isotype/SHM, and functional outputs.

In malaria, ABCs fluctuate in frequency: they peak around two weeks after symptom onset, decline within six weeks of treatment, and can remain elevated for up to six months. Many ABCs are non-switched, whereas class switched ABCs are enriched in previously exposed cohorts, consistent with antigen experienced memory (*9, 16*). Similar patterns occur in other diseases, including tuberculosis (*17*), hepatitis C (*18*), hantavirus infection (*19*), and COVID-19 (*20*), where CD11c+ T-bet+ ABCs emerge in response to strong inflammatory signals (*21*), to name a few. Beyond infection, ABCs are also a feature of vaccine responses: antigen-specific CD11c+ T-bet+ B cells expand transiently after influenza (*22*) and SARS-CoV-2 mRNA (*23*) vaccination, whereas an elevated baseline frequency of CD21low ABCs has been associated with poor vaccine responses (*24*). These observations highlight the common and dynamic nature of ABCs across immune contexts.

Current hypotheses propose multiple pathways for ABC differentiation, including from the splenic marginal zone (*21*) and germinal center derived memory B cells (*25–28*). BCR sequence data show that ABCs often carry somatic hypermutation at levels comparable to or lower than conventional memory B cells (*29*), suggesting germinal center experience. Yet subsets with lower SHM have also been identified in mice during infections such as LCMV, influenza or malaria (*10, 30, 31*), indicating possible germinal center independent origins.

The link between ABC emergence and impaired humoral responses has raised questions about their role in immunity and their functions remain debated across inflammatory contexts (*32*). ABCs may represent exhausted cells with reduced survival under chronic antigen load (*5*). Alternatively, they may act as plasma cell precursors (*33, 34*) or specialize in antigen presentation by upregulating MHC II and CD86 (*35*). These competing hypotheses underscore the complexity of ABC biology and the need for further research.

Evidence also indicates heterogeneity within ABCs (*36*), which may explain their diverse functions. However, the origins, relationships between potential subsets, and differentiation pathways remain poorly defined, contributing to variability and conflicting interpretations in the literature. Here, we combined high-dimensional flow cytometry, single-cell transcriptomics, V(D)J sequencing, and functional assays in human malaria and healthy donors to resolve ABC ontogeny and differentiation. We show that ABCs form a single CD11c/CD24/CD27 differentiation continuum, with distinct memory and extrafollicular origins converging on a shared late-stage phenotype and a stage-dependent shift in function that reconciles the conflicting roles previously attributed to ABCs.

## Results

### ABC-associated proteins are expressed by proliferating B cells during infection

B cell proliferation is crucial during infection, driving antigen-specific clonal expansion and the formation of effector and memory cells. Markers of proliferation can therefore indicate B cells contributing to the ongoing immune response. Several different proteins, including CD11c, T-bet, CXCR3, and FcRL5 combined with changes in CD21 and CD27 expression have been associated with ABCs. The diverse marker combinations used across studies, however, complicate direct comparisons and conclusions. Therefore, we adopted an unbiased approach, examining multiple markers associated with ABCs to better capture their complexity.

Mature B cells (CD19+CD10–CD38lo) from four individuals with recent malaria (**Table S1**) were assessed for T-Bet, CXCR3, CD11c, and FCRL5 expression in Ki67+ (recently dividing) and Ki67– (non-dividing) populations (**Fig. 1A-B**). Ki67+ B cells in *P. falciparum* infected patients showed consistently higher expression of these ABC-associated proteins compared to Ki67– cells (**Fig. 1C**). Using Boolean gating to identify all combinations of ABC-associated markers, we strikingly observed that 96.6% of Ki67+ B cells expressed one or a combination of ABC markers (**Fig. 1D**). The predominant subsets were quadruple-positive for T-Bet, FcRL5, CXCR3, and CD11c, and triple-positive for T-bet, CD11c, and FcRL5.

**Fig 1.**
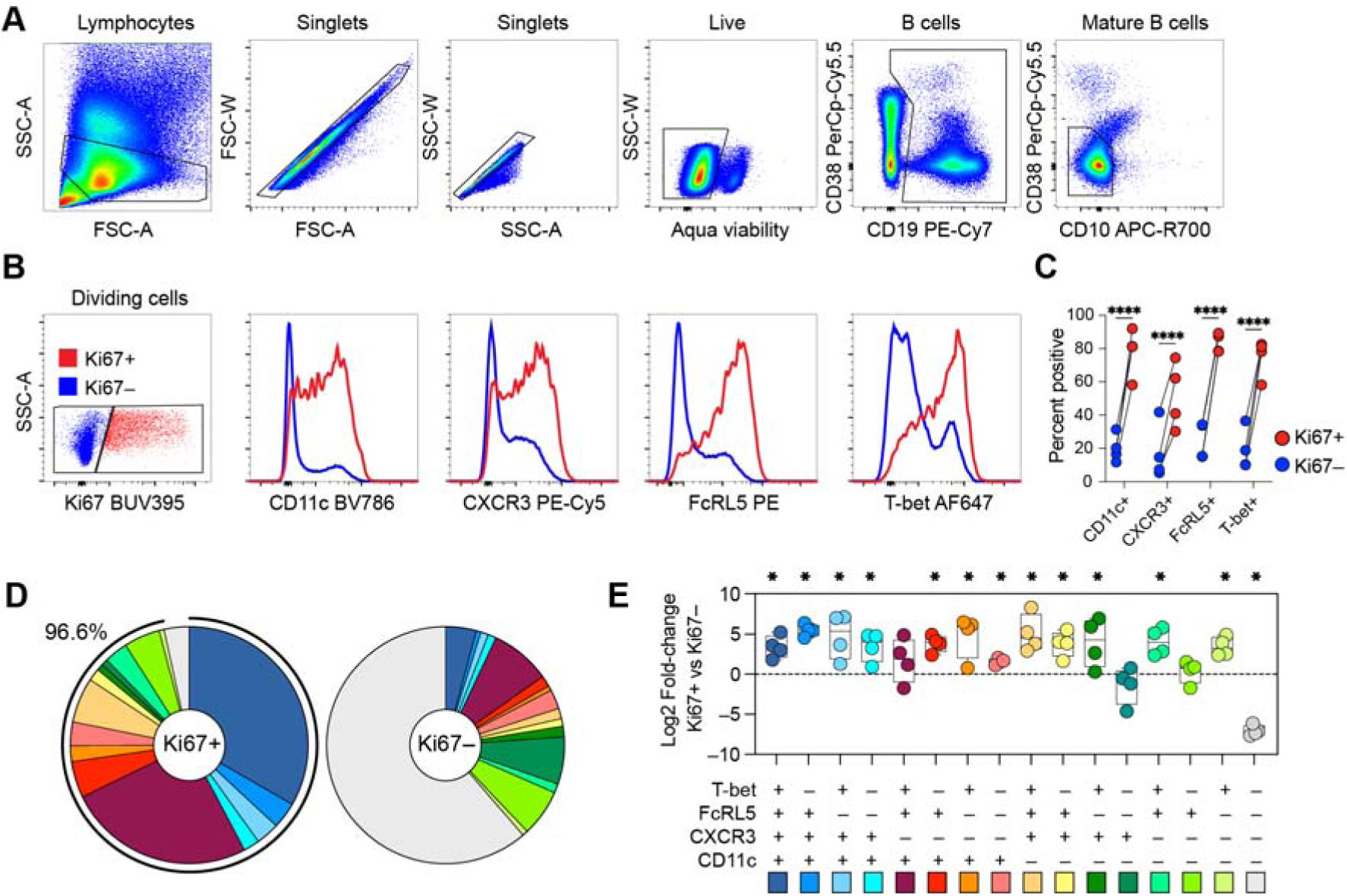
Proliferating circulating mature B cells express diverse marker combinations associated with ABCs. (**A**) Representative gating strategy of mature B cells and (**B**) CD11c, CXCR3, FcRL5, and T-bet expression among Ki67+(red) recently dividing and Ki67– (blue) non-dividing mature B cells. (**C**) Frequency of mature B cells positive for CD11c, T-bet, CXCR3, and FcRL5 among Ki67+ and Ki67– cells in patients with P. falciparum malaria. Statistics was evaluated using a paired two-way ANOV followed by Sidak’s post-hoc test, ****p<0.0001. (**D**) Donut plot showing the frequencies of populations expressing one or several atypical-associated markers among Ki67+ (left) or Ki67– (right) cells in P. falciparum malaria (n=4). (**E**) Log fold-change in the frequency of each marker combination between Ki67 and Ki67 mature B cells in individuals with P. falciparum malaria; values > 0 indicate relative enrichment (expansion) and values < 0 indicate depletion among proliferating (Ki67) cells. Statistics: ratio paired t-tests on Ki67+ versus Ki67– frequencies for each combination prior to fold-change calculation. *p < 0.05. Source Data for this Figure could not be made freely accessible, but have been deposited to the controlled-access repository Zenodo

For each marker combination, we compared its frequency between Ki67+ and Ki67– B cells, allowing us to determine which combinations had expanded (log fold-change > 0) or contracted (log fold-change < 0) in the proliferating compartment. (**Fig. 1E**). Overall, there was more expansion among B cells that expressed >1 ABC marker, potentially indicating that B cells with more ABC markers represent cells responding more strongly following stimulation or indicating more recent activation (**Fig. 1E**).

To assess if the heterogeneity and ABC marker pattern was specific to malaria, we also compared the results to previously published B cell responses characterised after Hantavirus infection (*19*) run in the same experiment (**Fig. S1**). We observed a similarly high level of ABC markers on Ki67+ B cells (**Fig. S1B**), with a high degree of heterogeneity among the markers (**Fig. S1B-C**). However, we also observed differences between malaria and Hantavirus infection, potentially indicating disease-specific ABC marker patterns (**Fig. S1D-E**).

In summary, ABC markers were strongly associated with B cell proliferation, as nearly all Ki67+ cells expressed at least one ABC-associated marker. The combination of markers was heterogeneous within malaria and also with hantavirus infection.

### Targeted CITE-Seq analysis reveals differentiation-driven heterogeneity of ABCs in malaria

To explore how ABC marker heterogeneity relates to B cell transcriptional profiles, we made use of a recently described targeted CITE-seq dataset, generated from PBMCs from four malaria patients sampled during acute infection, 10 days, and 1-year post-infection (*37*). Two patients had primary infections (malaria-naïve), and two had previously been exposed to malaria (**Table S1**). We specifically focused on the B cell data and extracted and re-clustered the cells using the Seurat pipeline, with clustering done based on the mRNA data (**Fig. 2A and Fig. S2A-C**).

**Fig 2.**
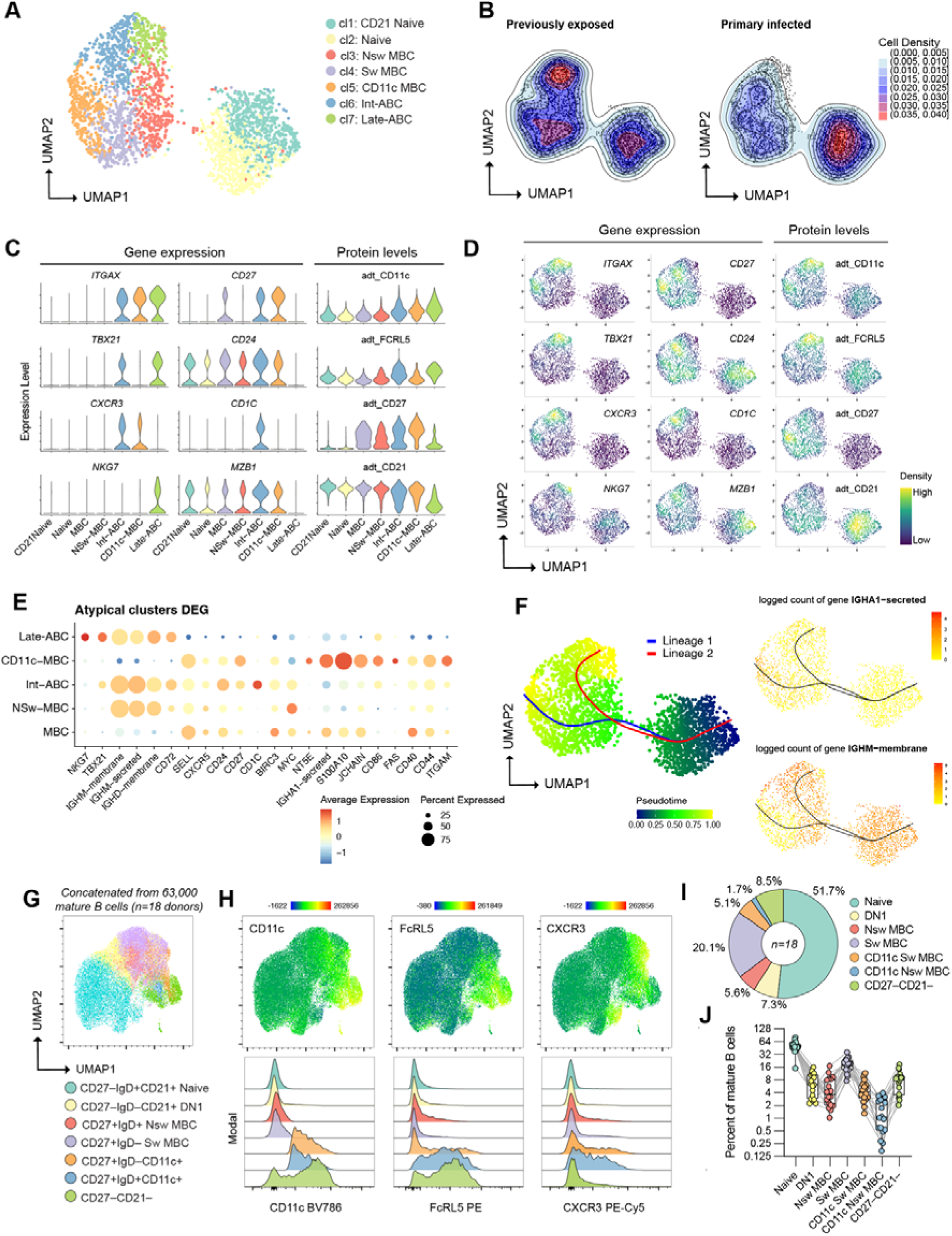
Targeted CITE-Seq analysis of circulating B cells in malaria patients. (**A**) UMAP indicating B cell clusters done on 2601 cells from 12 pooled samples (4 donors and 3 time-points after malaria). (**B**) Comparison of cell densities between previously exposed (left, n=2, 1631 cells) and primary infected (right, n=2, 970 cells) individuals. (**C**) Violin plots displaying gene expression distributions of atypical B cells (ABC) markers and associated surface proteins (ADT). (**D**) Density plots indicating gene and protein (ADT) expression distributions of selected ABC-associated markers. (**E**) Dot plot showing the top 22 differentially expressed genes across memory and ABC clusters. (**F**) UMAP colored by Slingshot pseudotime, with smoothed curves representing two lineages and IgA/IgM expression across pseudotime. (**G**) UMAP of mature B cells concatenated from 18 donors with acute malaria colored with cell subsets identified by flow cytometry corresponding to scRNAseq clusters. (**H**) CD11c, FcRL5, and CXCR3 protein intensity (top panel) and expression in the different cell populations (bottom panel). (**I-J**) Overall (**I**) and individual donor (**J**) frequency of each population among total mature B cells. Source Data for this Figure could not be made freely accessible, but have been deposited to the controlled-access repository Zenodo

The clustering identified seven distinct clusters, with most evenly distributed between primary and previously infected individuals, except cluster 7, which was enriched in cells from previously infected patients, potentially indicating a memory origin (**Fig. 2A-B, Fig S2D-F**). Investigating the expression pattern of each cluster allowed us to identify two naïve clusters (cluster 1 and 2), and five memory-associated clusters (3–7) (**Fig. S2G**). Specifically analyzing the clusters for ABC markers identified one non-switch memory cluster (cl 3), one switch memory cluster (cl 4) and three clusters (cl 5-7) expressing different patterns of ABC-associated markers (**Fig. 2C-D and Fig. S2H-I and Fig. S3A**). Cluster 5 expressed *ITGAX* (CD11c) and *CXCR3* but lacked *TBX21* (T-bet), except transiently on day 10, pointing to them as potential ABC precursors (**Fig. 2C-D and Fig. S3B**). Consistent with this, they also retained their expression of *CD27* and *SELL* (CD62L) (**Fig. 2C-E**). Cluster 6 co-expressed *ITGAX, TBX21, CXCR3,* adt_FCRL5 (protein levels), *CD1c*, and similar expression of *IGHM* as non-switch memory cells (cl 3), while cluster 7 was associated with high levels of *ITGAX, TBX21* and *NKG7* and low levels of *CXCR3*, CD21 and CD27 protein levels and *CD27* gene expression (**Fig. 2C-D**). These features along with an upregulation of FCRL5 protein levels suggested that cluster 7 represents a late stage of differentiation. Based on these data we denoted cluster 5 as CD11c+ memory B cells (Early-ABC), cluster 6 as intermediate atypical memory B cells (Int-ABC), and cluster 7 as late-stage atypical B cells (Late-ABC).

We next analyzed the top 22 differentially expressed genes (DEG) distinguishing the five memory clusters (**Fig. 2E**). Late-ABC and Int-ABC showed enrichment in non-switched BCR isotypes, while Early ABC were enriched in switched isotypes, particularly IgA. Notably, CD24 and CD27 expression patterns differed: Int-ABC exhibited high CD24 and low CD27, while Early ABC showed the opposite. Both markers were downregulated in Late-ABC, consistent with their potential late stage of differentiation (**Fig. 2E**).

To assess these possible connections between populations, we performed a pseudotime trajectory analysis and identified two lineages (**Fig. 2F and Fig. S3C**). They first progressed from naïve B cells through non-switched MBCs and Int-ABC to Late-ABC, reflecting a differentiation pathway toward progressively later ABCs. The second passed through switched MBCs and culminated in Early ABC, indicating a distinct route largely influenced by isotype switching, particularly IgA. This divergence may reflect functional specialization shaped by isotype-specific roles in immunity or distinct developmental origins driven by infection-induced cues. Whether these pathways serve unique immune functions or represent alternative differentiation trajectories is unclear. Importantly, these trajectories reflect relationships between transcriptional states and do not imply a single linear differentiation pathway or direct transitions between all populations, as Slingshot relies on local structure in the low-dimensional embedding and cannot reliably infer relationships between non-adjacent populations.

While the CITE-seq dataset enabled high-resolution identification of ABC states and their differentiation trajectories in a longitudinal setting, we next used flow cytometry in a larger acute malaria cohort to validate these populations at the protein level and assess their frequency across individuals. We re-analysed an acute-malaria dataset of 18 patients (**Table S1**); all processed in a single experiment to reduce batch variation (*16*). From each donor we concatenated 3,500 mature B cells (excluding CD38high CD20low plasma cells and CD10+ immature B cells; total 63,000) and applied UMAP for dimensionality reduction. CD27, IgD, CD21, and CD11c were used as defining markers to delineate populations corresponding to the clusters (**Fig. 2G**). The two naïve clusters (1 and 2) from the scRNA-seq analysis were represented as a single CD27–IgD+CD21+ naïve population. Switched and non-switched MBCs were defined by CD27 and IgD, and CD11c-MBC (Early-ABC) and Int-ABC were further resolved by gating for CD11c within the switched and non-switched MBC compartments, respectively. Late ABCs were identified by low expression of CD21 and CD27 (**Fig. 2G**). Protein levels of CD11c, FCRL5, and CXCR3 aligned with the cluster expression profiles (**Fig. 2H**). Overall frequencies are shown in **Fig. 2I**, with per-donor variation in **Fig. 2J**. Protein-level analyses confirmed the RNA-seq-derived clusters in a larger patient cohort, highlighting their consistency across method and patients.

Although not identified as a cluster in the scRNAseq analysis, we included a CD27–IgD–CD21+ population, previously identified as DN1 cells and proposed as precursors for switch MBCs (*38*). Other studies have also suggested that the CD21–CD27– Late-ABC cluster can be further subdivided into IgD–CD11c+ DN2 cells, suggested to be antibody-secreting cell (ASC) precursors, and IgD–CD11c– DN3 cells, suggested to be precursors of DN2 cells (*38*). However, such gating which only includes switched cells excludes approximately 35% of Late-ABCs (**Fig. S3D**), potentially limiting our understanding of ABC differentiation and function in malaria where non-switched ABCs were shown to greatly expand after infection (*16*).

The analysis of longitudinal samples can help identify specific subsets responding during and after acute malaria infection. We therefore extended our analysis from Fig. 2G-J to also include time-points up to one year after acute infection and further analyzed the samples with regards to those with primary infection and those with previous malaria exposure (**Table S1**), to indicate potential impact of pre-existing memory (**Fig. S4**). There were no differences in Naïve and switch MBC frequencies between groups or over time, while DN1 cells had minor changes over time. Non-switch MBCs were more frequent in primary infected individuals,

but there was no association with time (**Fig. S4A**). CD11c+ switch and non-switch MBCs and CD27–CD21– B cells all had significant associations with time, with a peak expansion by day 10 after acute malaria, which then contracted over time. Both CD11c+ switch MBC and CD27–CD21– B cells were significantly more expanded in previously exposed individuals while the expansion of CD11c+ non-switch MBCs was independent of memory (**Fig. S4A**). Further subsetting the CD27–CD21– B cells into IgD+ and IgD– DN2 and DN3 cells showed a significant time-effect on DN2 cells but not DN3 cells. The expansion of IgD– DN2 cells was primarily observed in previously exposed individuals while the expansion of IgD+ DN2 cells was independent of prior malaria exposure (**Fig. S4B**). We also assessed time-dependent changes in the clusters from the CITE-seq data with an early expansion of CD11c-MBC and Int-ABC clusters and a slightly delayed expansion of Late-ABCs (**Fig. S4C**).

Together, the targeted CITE-seq dataset provided a longitudinal, high-dimensional framework for defining ABC states and their differentiation trajectories, whereas flow cytometry enabled validation of these populations at the protein level and assessment of their distribution across a larger patient cohort over time and with regards to pre-existing memory.

### High-resolution transcriptomic and V(D)J sequencing of CD11c+ B Cells reveals additional intermediate ABC states with different plasma cell differentiation potential

Our flow cytometry and CITE-seq analysis identified CD11c as the minimum common feature of potential ABC lineages derived from memory B cells. To gain a more detailed understanding of CD11c+ B cell differentiation after malaria, we performed 10X scRNA-seq with paired V(D)J sequencing on sorted CD11c+ B cells, 10 days after treatment with artemether/lumefantrine.

We identified five distinct clusters within the CD11c+ B cell compartment (**Fig. 3A and Fig. S5A**), four of which expressed multiple ABC markers and one small cluster with very few cells expressing CD69, potentially representing recently activated memory B cells or cells migrating into tissues (**Fig. 3B and Fig. S5A**) (*39*). Consistent with the sorting strategy, all clusters expressed *ITGAX* (CD11c). They also expressed transcription factor *ZEB2*, which was recently proposed as a key driver for CD11c expression in B cells (**Fig. 3B**) (*40, 41*). Based on the flow cytometry data and CITE-Seq definitions determined in Figure 2, we identified Early ABC cells expressing high levels of *CD24*, *CD27* and relatively low levels of *ITGAX*, *CXCR3*, and *FCRL5* and no detectable *TBX21* (T-bet) or *NKG7* (**Fig. 3B and Fig. S5B**). We also identified Late-ABCs, defined by high *TBX21*, *ITGAX*, *FCRL5*, and *NKG7* expression but lacking *CXCR3*. Contrasting with the targeted CITE-Seq dataset, we identified two intermediate clusters instead of one; Int-ABC1 could be distinguished as *CD24–CD27+* and Int-ABC2, distinguished as *CD24+CD27–* (**Fig. 3B and Fig. S5B**).

**Fig 3.**
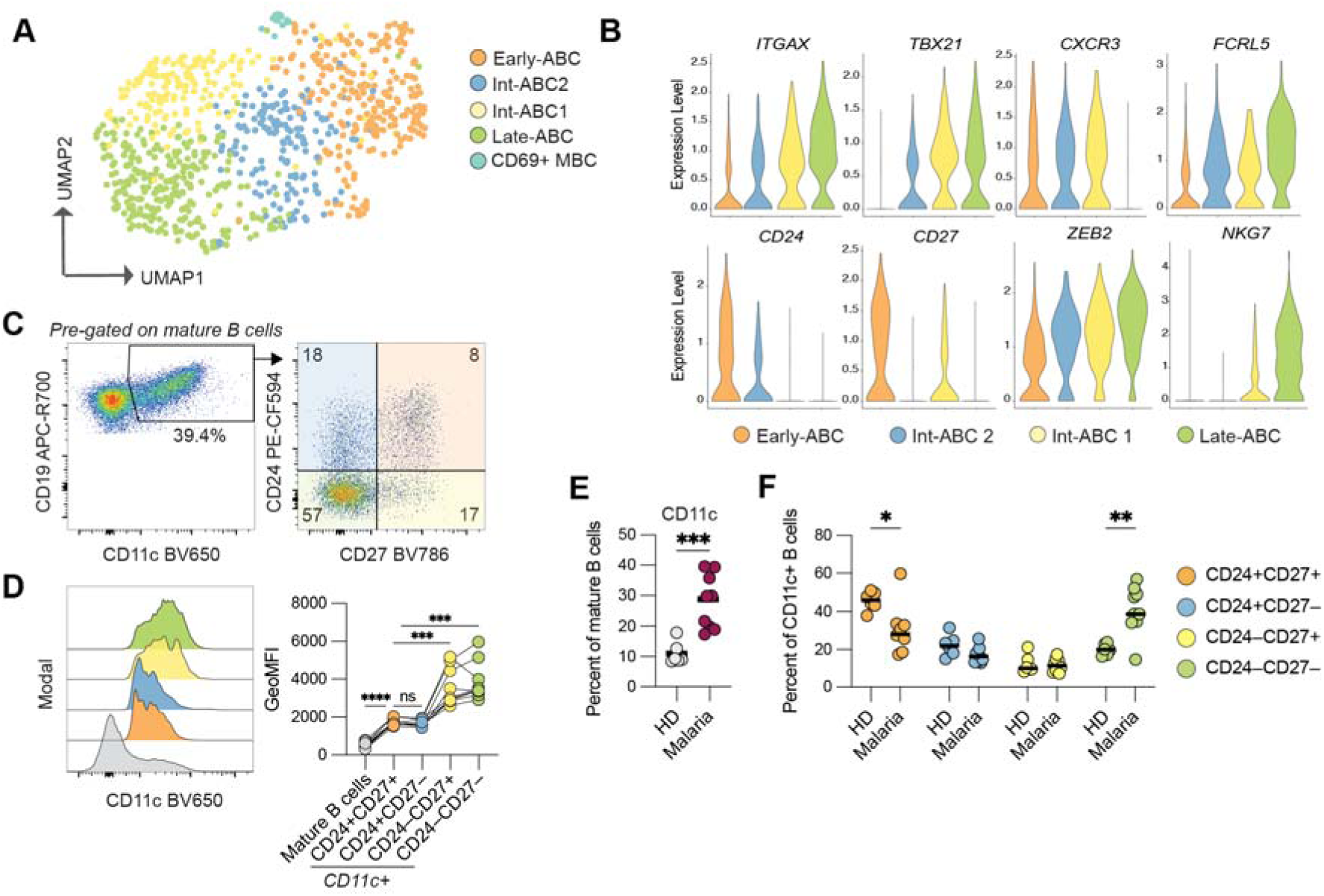
scRNA-Seq Analysis of sorted CD11c+ B Cells at two weeks Post-Malaria. (**A**) UMAP showing distinct CD11c+ B cell subsets in a malaria patient two weeks post diagnosis. (**B**) Violin plots displaying gene expression distributions of genes associated with atypical B cells. (**C**) Representative gating for CD19+CD11c+ followed by CD24 and CD27 among mature B cells two weeks after malaria diagnosis. (**D**) Representative histogram (left) and comparisons (right) of CD11c geometric mean intensity of total mature B cells (grey), and CD24+CD27+ (orange, Early-ABC), CD24+CD27– (blue, Int-ABC2), CD24–CD27+ (yellow, Int-ABC1), and CD24–CD27– (green, Late-ABC) CD11c+ mature B cells for malaria donors (n=9). Statistical comparisons were done using matched pair one-way ANOVA. (**E**) Frequency of CD19+CD11c+ cells among mature B cells in healthy donors (grey circles, n=6) and malaria donors (maroon circles, n=9), unpaired student’s t-test with Welch’s correction. (**F**) Frequency of cells expressing different combinations of CD24 and CD27 among CD19+CD11c+ mature B cells for healthy donors (n=6) and malaria donors (n=9). Statistical comparisons were done using two-way ANOVA followed by Sidak’s posttest. *p<0.05, **p<0.01. Source Data for this Figure could not be made freely accessible, but have been deposited to the controlled-access repository Zenodo.

To assess whether intermediate ABC stages could also be resolved at the protein level, we analyzed CD11c+ mature B cells from malaria patients 10 days post-treatment and compared them with healthy controls (**Table S1**). CD24 and CD27 distinguished four CD11c+ B cell subsets (**Fig. 3C**), with protein levels overlapping the *ITGAX* gene expression patterns (**Fig. 3D**). CD11c+ B cells were more frequent in malaria patients (**Fig. 3E**). Among them, intermediate subsets were comparable between groups (∼12% Int-ABC1, ∼20% Int-ABC2), while early ABCs (CD24+CD27+) were enriched in healthy donors and late ABCs (CD24–CD27–) in malaria patients (**Fig. 3F**).

In summary, CD11c B cells comprised early, late, and two intermediate ABC subsets defined by CD24 and CD27 expression and transcriptional profiles. Their distribution varied by infection status, with early ABCs enriched in healthy donors and late ABCs in malaria patients.

### Distinct isotype profiles and origins of intermediate ABC subsets

To further understand the ABC differentiation process, we analyzed V(D)J sequencing data from the dataset described in Fig. 3. We found that all clusters were predominantly composed of B cells with a non-switched *IGHM* BCR, with both intermediate clusters (Int-ABC1: 83.9%, Int-ABC2: 95%) to a larger extent than Early-ABCs (68%) and the Late-ABCs (79%). BCRs were detected from a wide range of V-genes with relatively minor differences between the clusters (**Fig. S6A-B**). Consistent with the transcriptomic data, the Early-ABC cluster contained the largest number of B cells expressing an *IGHA* isotype, with a small proportion also present in the Int-ABC1 cluster. Additionally, Int-ABC1 also exhibited a higher proportion of switched isotypes (16%) compared to Int-ABC2 (5%) (**Fig. 4A**).

**Fig 4:**
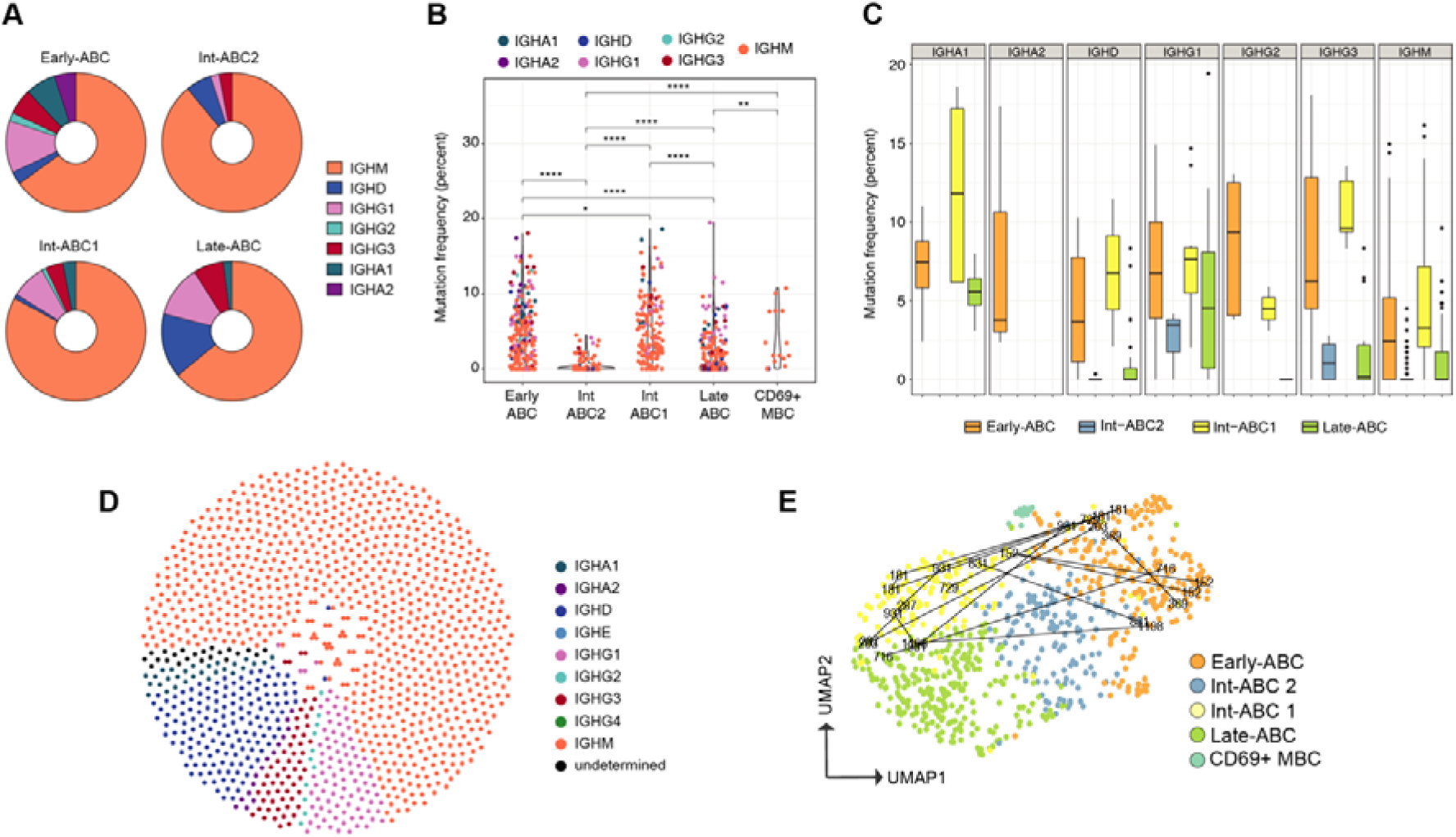
Isotype Usage Patterns Across Atypical B Cell Subsets Revealed by V(D)J Sequencing. (**A**) Isotype use among atypical B cell (ABC) clusters (**B**) V-gene mutation frequency among ABC clusters. Statistical analyses were done using unpaired Wilcoxon test with *p<0.05, **p<0.01, ***p<0.001, ****p<0.0001. (**C**) Mutation frequency among isotypes in each cluster. (**D**) Honeycomb B cell receptor clonal distribution with each dot representing one cell and cells from the same lineage clustered together. (**E**) UMAP of gene expression data with lines indicating clonally related B cells. Source Data for this Figure could not be made freely accessible, but have been deposited to the controlled-access repository Zenodo.

We next examined the heavy chain V-gene sequences of each cluster for somatic hypermutation. Int-ABC2 (average: 0.4%) displayed significantly lower mutation frequency compared with Int-ABC1 (average: 5.4%) which was similar to Early-ABC (average: 4.5%), while Late-ABC (average: 1.6%) exhibited an intermediate mutation frequency between Int-ABC2 and Int-ABC1 (**Fig. 4B**). Detailed examination of mutation frequencies by isotype confirmed this pattern, with Int-ABC1 consistently showing similar mutation levels to Early-ABC across isotypes, including for non-switched cells (**Fig. 4C**).

To investigate inter cluster relationships, we examined clonal lineages. Few lineages were detected (**Fig. 4D**), in line with CD11c based sorting rather than antigen specific enrichment. Among the lineages detected, several bridged Early-ABC and Int ABC1 (**Fig. 4E**), and some extended across additional clusters and isotypes (**Suppl. Fig. 6C).**

In summary, the two intermediate subsets showed distinct BCR profiles, potentially indicating different origins. Int-ABC1 displayed higher mutation frequencies and greater isotype diversity, consistent with having gone through a germinal center (GC) reaction. Int-ABC2 was enriched for IGHM with low mutation frequencies, indicating origins from naïve or non-switched memory cells generated in extrafollicular reactions. Late ABCs exhibited intermediate levels of switched isotypes and somatic hypermutation, potentially indicating contributions from both intermediate subsets. Importantly, Early ABC align with the mutated, often class-switched Int-ABC1 axis, whereas Int-ABC2 represents a distinct population with low somatic hypermutation, most consistent with arising from a non-switched, antigen-experienced precursor pool.

### CD24 and CD27 downregulation during ABC differentiation is isotype-dependent

To test whether the observed clusters reflect a continuum of differentiation, we analyzed CD24 and CD27 dynamics, key markers distinguishing the ABC subsets, *in vitro* and *ex vivo*. Using a previously described *in vitro* ABC differentiation system (*42*), we tracked stepwise changes in marker expression in sorted mature B cells (**Fig. S7A and S7B**). This model recapitulated hallmark features of ABC differentiation, as confirmed by upregulation of CD11c, CD1c, T-bet, CXCR3, and NKG7 protein levels (**Fig. 5A and Fig. S7C**). Using Boolean gating, we could further see that differentiation was associated with varied co-expression of ABC-associated markers with few cells (<10%) not expressing any ABC-associated proteins at 2 and 4 days of stimulation, reminiscent of the pattern observed during acute malaria and hantavirus infection (**Fig. S7D**).

**Fig 5:**
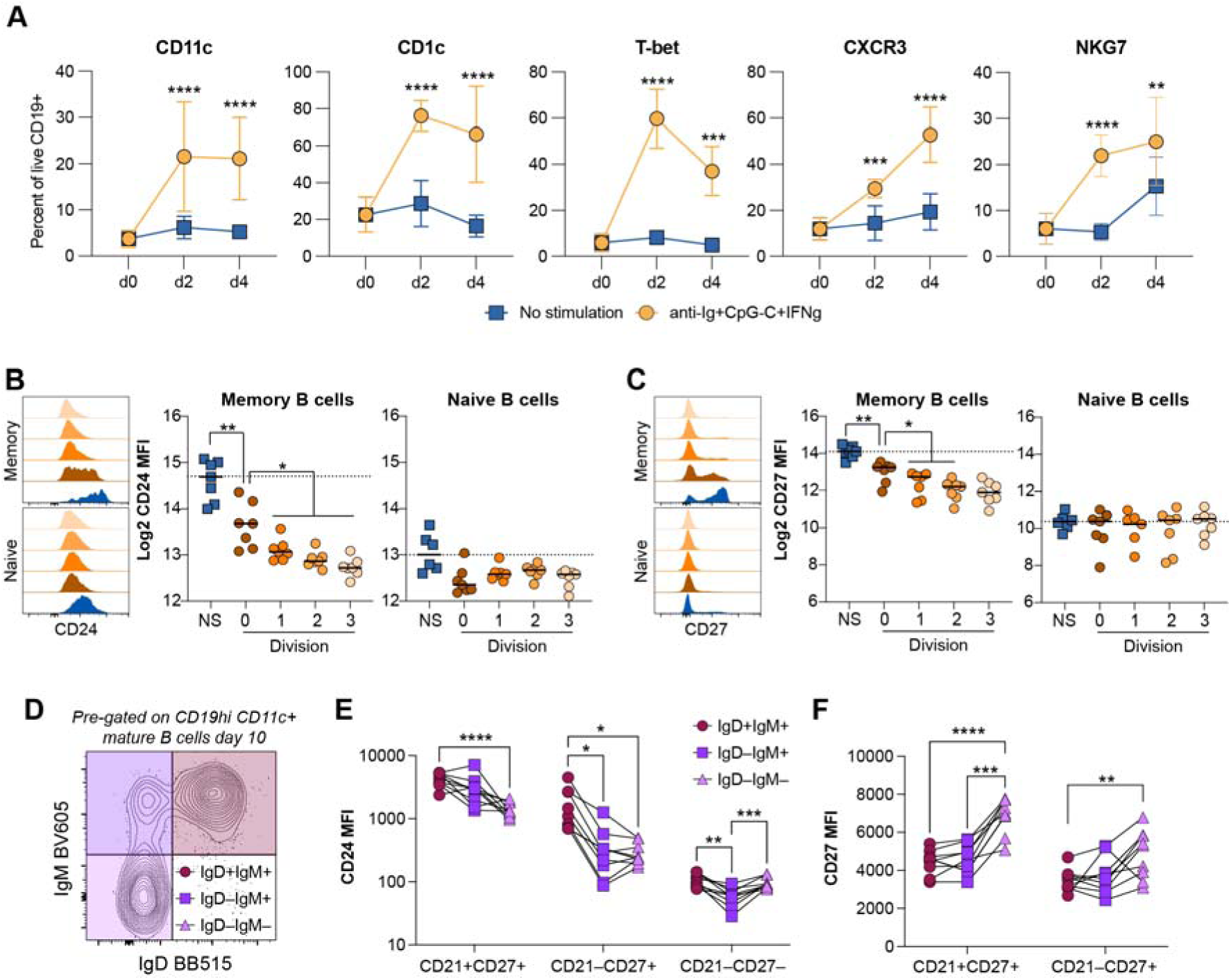
In Vitro and Ex vivo analysis reveals CD24 and CD27 dynamics in ABC differentiation. (**A**) Longitudinal analysis of surface expression of CD11c, CD1c, T-bet, CXCR3, and NKG7 on non-stimulated (media, blue boxes) or stimulated (anti-Ig, CpG-C, IFNγ, yellow circles) sorted human total B cells from healthy blood donors (n=4) at day 0, 2, and 4. Error bars indicate the standard deviation. Statistical analyses were done using matched pair two-way ANOVA with Dunnet’s posttest. **p<0.01, ***p<0.001, ****p<0.0001. (**B-C**) Expression of CD24 (**B**) and CD27 (**C**) in non-stimulated (NS, blue boxes) or stimulated (anti-Ig+CpG-C+IFNγ, circles with orange shades) sorted memory and naïve B cells at day 4 of culture. Stimulated cells were gated by number of cell divisions (**Suppl. Fig 7A-B**). Statistical analyses were done using repeated measures one-way ANOVA followed by Sidak’s post-hoc test. *p<0.05, **p<0.01. (**D**) Representative gating strategy of IgD+IgM+, IgD–IgM+, and IgD–IgM– CD19hiCD11c+ B cells. (**E-F**) Median fluorescent intensity (MFI) of CD24 (**E**) and CD27 (**F**) among CD21+CD27+ resting memory, CD21–CD27+ activated memory, and CD21–CD27– double negative B cells at day 10 after acute malaria (n=9). Statistical analyses were done using two-way ANOVA followed by Tukey’s post-hoc test. *p<0.05, **p<0.01, ***p<0.001, ****p<0.0001. Source Data for this Figure could not be made freely accessible, but have been deposited to the controlled-access repository Zenodo.

We next stimulated sorted memory and naïve B cells and tracked CD24 by flow cytometry. Memory B cells showed marked CD24 downregulation after stimulation, with further decreases accompanying cell division. In contrast, naïve B cells showed no significant change, potentially reflecting the lower CD24 starting levels compared with memory B cells and that the levels declined in culture independent on stimuli (**Fig. 5B**). Similarly, CD27 expression decreased in memory B cells upon stimulation and after division, and as expected remained absent in naïve B cells (**Fig. 5C**). These findings suggest that the CD24+ Int-ABC2 subset likely originates from marginal zone-like non-switched memory B cells rather than naïve B cells, and that memory B cells progressively lose CD24 and CD27 as they differentiate toward a late-stage ABC phenotype.

To confirm that the *in vitro* data reflected the *in vivo* situation, we examined CD24, CD27, and CD21 levels during ABC differentiation in PBMCs from individuals with acute malaria. CD11c+ mature B cells were stratified by IgM and IgD expression to represent GC-derived class switched (IgD–IgM–), GC-derived non-switched (IgD–IgM+) and marginal zone-like non-switched (IgD+IgM+) B cells (**Fig. 5D**) (*43*). These were then gated as starting their differentiation (CD21+CD27+), ongoing differentiation (CD21–CD27+), and late-stage differentiation (CD21–CD27–). Naïve (CD21+CD27–) B cells were not included in the analysis. CD24 median fluorescent intensity (MFI) was assessed and declined with differentiation, consistent with the *in vitro* data (**Fig. 5E** and **Fig. S7E**). The marginal zone-like B cells (IgD+IgM+) retained significantly higher CD24 levels than the switched and non-switched memory cells (IgD–IgM– and IgD–IgM+) during ongoing differentiation, while the GC-derived cells had retained similar levels (**Fig. 5E**). For CD27, we could not assess levels at the late stage, as those are defined as CD27–. However, consistent with the *in vitro* data, CD27 levels declined with B cell activation and differentiation (**Fig. 5F** and **Fig. S7E**). Additionally, both at the start of differentiation and during ongoing differentiation, the levels were significantly higher for the switched memory cells.

These findings support that the intermediate ABC clusters represent transition states of cells with different origins, where Int-ABC1 originates from GC-derived switched (IgD–IgM–) and IgD–IgM+ B cells while Int-ABC2 originates from IgD+IgM+ marginal zone-like B cells, that both eventually converge into Late-ABC.

### Conserved heterogeneity of ABC subsets in secondary lymphoid tissue of healthy donors

Since B cell activation and maturation primarily occur in secondary lymphoid organs, and intermediate subsets were indicated to have originated from GC or marginal zone-like B cells following extrafollicular B cell activation, we assessed the presence and characteristics of early, intermediate, and late ABCs in tonsils and paired peripheral blood samples.

To compare the overall distribution and phenotype of ABCs across blood and tonsils, we concatenated total B cells from four healthy donors and visualized them by UMAP (**Fig. 6A**). We gated cell subsets as done for malaria samples (**Fig. 2G**) with the addition of gates for GC B cells, transitional cells, and plasma cells (**Fig. S8A-B**). Overall, B cell subsets were similar with the clear exception of GC cells that were present in tonsils but not PBMCs and transitional cells which were abundant among PBMCs (**Fig. 6B**). Both tissues contained early, intermediate, and late ABC subsets with broadly similar distribution (**Fig. 6B–C**). CD11c was expressed in both PBMC and tonsil B cells (**Fig. 6D**), and the proportion of CD11c+ mature B cells was largely comparable, with a non significant trend toward overall lower frequencies in tonsils (**Fig. 6E**). Almost all CD11c+ B cells were present in the mature B cell compartment (i.e., CD19+ B cells excluding transitional, plasma, and germinal-center cells) (**Fig. 6F**). Notably CD11c+ B cells were not observed within the GC B cell gate, suggesting they may be excluded from ongoing GC reactions. CD11c expression was however highly enriched among Ki67+ B cells, indicating recent activation and cell division (**Fig S8C**).

**Fig 6:**
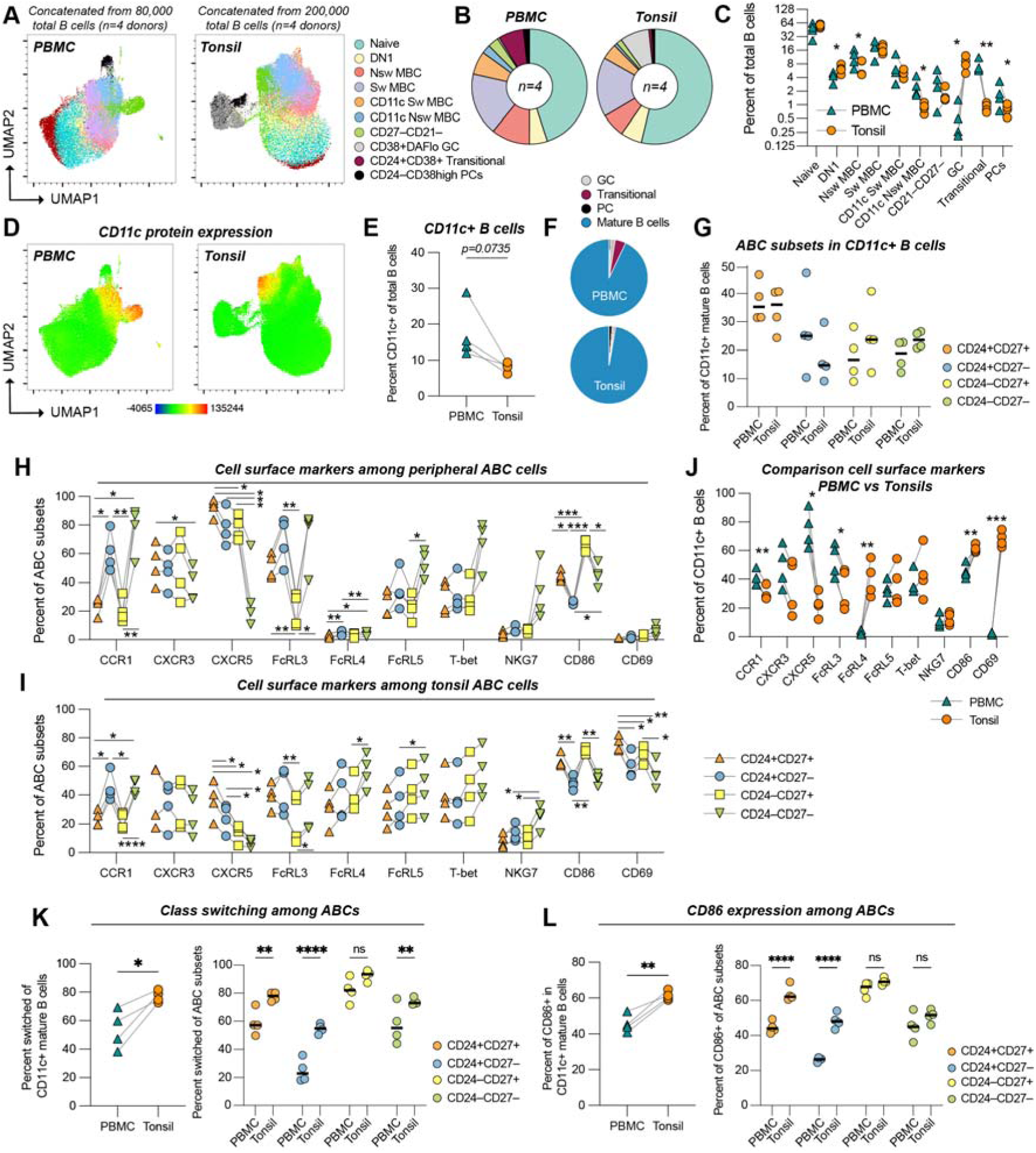
Blood and tonsil have similar distribution of early, intermediate and late ABCs but they display differences in ABC switching and costimulatory molecules. **(A)** UMAP projection of total B cells analyzed in PBMC and tonsil samples, concatenated from four healthy donors. Cell clusters are colored by subsets identified by flow cytometry. **(B)** Overall proportion of each identified subset within each tissue. **(C)** Comparison of subset frequencies of total B cells between PBMC and tonsil. Statistical analyses were done using matched pair two-way ANOVA on log2 transformed data. **(D)** UMAP illustrating CD11c protein intensity. **(E)** Frequency of CD11c+ mature B cells in paired PBMC and tonsil samples (n = 4). Statistical analysis with a paired t-test. **(F)** Distribution of CD11c+ B cells across germinal center (GC), transitional, plasma cell (PC), and mature B cell compartments. **(G)** Frequencies of CD24+CD27+ (Early-ABC), CD24+CD27– (Int-ABC2), CD24–CD27+ (Int-ABC1), and CD24–CD27– (Late-ABC) subsets within CD11c+ B cells in PBMC and tonsil. Statistical analyses with matched pair two-way ANOVA. All p>0.05. **(H)** Frequency of cells expressing indicated markers in CD24/CD27-defined ABC subsets in PBMCs **(H)** and tonsils **(I)**. Statistical analyses by matched pair two-way ANOVA. **(J)** Frequency of marker expression among CD11c+ mature B cells from PBMC (blue triangle) and tonsils (orange circle). Statistical analyses as in (C). **(K)** Frequency of switched (IgD–IgM–) cells among CD11c+ B cells, and within each CD24/CD27-defined subset, in PBMC and tonsil. Statistical analyses with paired t-test (left) or matched pair two-way ANOVA (right). Ns=p>0.05, *p<0.05, **p<0.01, ***p<0.001, ****p<0.0001. **(L)** Frequency of CD86+ B cells among CD11c+ B cells (left), and of each CD24/CD27-defined subset, in PBMC and tonsil (right). Statistical analyses as in (K). Source Data for this Figure could not be made freely accessible, but have been deposited to the controlled-access repository Zenodo.

We next stratified CD11c+ B cells by CD24 and CD27, to resolve early, intermediate, and late ABC subsets, and observed similar distributions between blood and tonsil (**Fig. 6G**). We further assessed protein markers associated with migration (CCR1, CXCR3, CXCR5, CD69), differentiation (T-bet), and function/activation (FcRL3, FcRL4, FcRL5, NKG7, CD86) between ABC subsets in PBMCs (**Fig. 6H**) and tonsils (**Fig. 6I**). The ABC subsets displayed overall similar patterns between the tissues, although many of the proteins were expressed to different extents (**Fig. 6J**). CCR1, CXCR5, and FcRL3 were more frequently expressed by PBMC ABCs, while FcRL4, CD86, and CD69 were more frequently expressed by tonsil ABCs (**Fig 6J**). BCR isotype analysis further revealed more switched (IgD–IgM–) CD11c+ B cells in tonsils compared to PBMCs, and this enrichment was observed in all subsets except Int-ABC1 (CD24–CD27+) where almost all cells were switched in both PBMC and tonsil samples (**Fig. 6K**). Increased class-switching was a general feature of CD11c+ B cells (**Fig. S8D**) and together with recent activation and cell division, could indicate more extensive interactions with cognate CD4+ T cells. We therefore assessed levels of the costimulatory molecule CD86 and observed an evident increase in both the overall CD11c+ population compared with CD11c– mature B cells (**Fig. S8E**) and within ABC subsets, closely mimicking the pattern for switched cells (**Fig. 6L**).

Together, these results indicate that the fundamental architecture of ABC subsets is conserved between blood and tonsil and can be captured by CD11c, CD24, and CD27, although ABC cell behavior could potentially differ as several proteins associated with cell function and migration were expressed to different extents between tissues.

### ABC subsets display stage-dependent functional profiles at the transcriptomic and ex vivo levels

Having established that ABC subsets were conserved across blood and tonsil, we next characterised their functional properties. Using the 10X scRNA-seq dataset, we performed ingenuity pathway analysis (IPA) canonical pathway enrichment analysis across Early-, Intermediate-, and Late-ABC clusters (**Fig. 7A**). Pathways related to phagosome formation and Fcγ receptor-dependent phagocytosis were progressively enriched in Intermediate- and Late-ABCs, whereas the Th1 pathway and MHC class II antigen presentation were more prominent in earlier clusters. To complement this, per-cell module scoring using UCell was applied to quantify transcriptional programs related to phagocytosis, cytokine production, antigen presentation, and antibody secretion across the four CD11c+ clusters (**Fig. 7B-C**). This revealed a striking functional dichotomy along the differentiation axis: Early-ABCs showed the highest scores for cytokine production and antibody secretion, whereas Late-ABCs displayed the highest scores for phagocytosis and antigen presentation. Intermediate subsets occupied transitional positions, with Int-ABC1 shifted toward phagocytosis and antigen presentation, and Int-ABC2 having partial cytokine and antibody secretion signatures.

**Figure 7.**
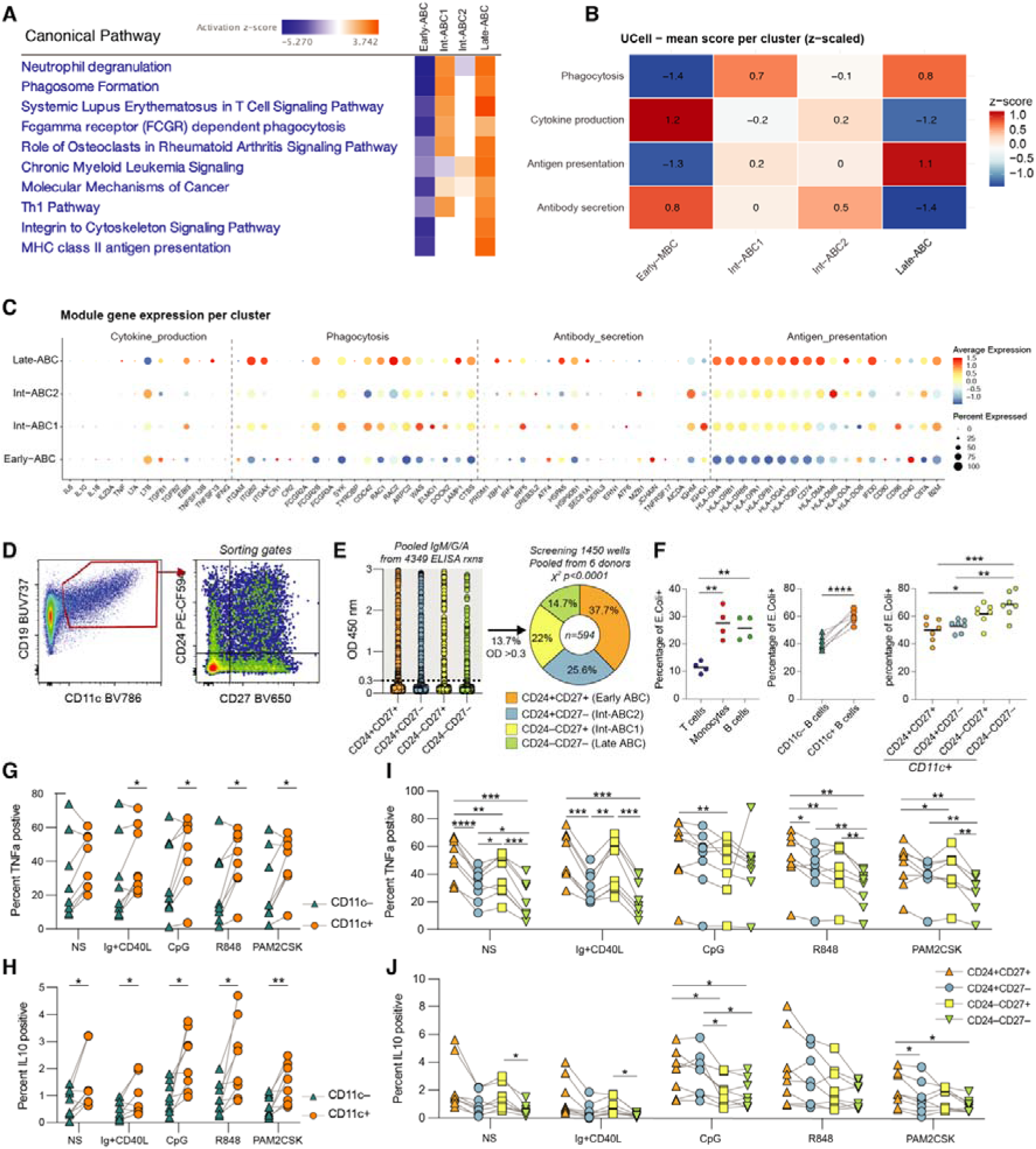
Functional characterization of ABC subsets at the transcriptional and protein level. **(A)** Ingenuity Pathway Analysis (IPA) of canonical pathways differentially enriched across ABC developmental stages (Early-, Int-, and Late-ABC) identified from previous scRNA-seq data. Heatmap displays activation z-scores. **(B)** UCell module scores (z-scaled) for functional gene sets (phagocytosis, cytokine production, antigen presentation, and antibody secretion) computed per cluster across CD11c+ B cells, intermediate ABCs (Early-ABC, Int-ABC1, Int-ABC2), and terminally differentiated ABCs (Ter-ABC). **(C)** Dot plot showing average expression and percentage of cells expressing individual module genes across clusters. Dot size reflects the fraction of expressing cells; color intensity reflects average expression level. **(D)** Gating for CD11c+ mature B cells followed by CD24 and CD27 for single B cell culture. **(E)** (Left) Pooled IgM, IgG, IgA antibody levels from individual sorted wells (n=1,450 wells and 4,349 data-points) as determined by ELISA. (Right) Distribution of ELISA-positive (OD > 0.3) wells among sorted B cell populations. Statistical evaluation was done with a Chi-square test, p<0.0001. **(F)** B cells isolated from PBMCs of 7 healthy donors were co-cultured with fluorescently labeled *E. coli* fragments and fluorescent particle association was assessed by flow cytometry. The percentage of *E. coli*-positive cells is shown for total B cells, T cells, monocytes, and B cell subsets stratified by CD11c and CD24/CD27 expression. **(G**–**J)** B cells isolated from PBMCs of 8 healthy donors were stimulated for 24 h with anti-Ig/CD40L, CpG, R848, or PAM2CSK4, followed by 4 h of PMA/ionomycin stimulation in the presence of brefeldin A and monensin. Intracellular cytokine staining was performed by flow cytometry. Percentage of TNFα+ **(G)** or IL-10+ **(H)** cells is shown for CD11c versus CD11c+ B cells. Percentage of TNFα+ **(I)** or IL-10+ **(J)** cells is shown for CD24/CD27-defined subsets within CD11c B cells. The 8 donors were processed in two independent experiments (two donor batches; n = 5 and n=3), with both TNFα and IL-10 measured in every experiment. Each dot represents one donor. Statistical comparisons by paired t-test or repeated-measures ANOVA with post-hoc correction; *p<0.05, **p<0.01, ***p<0.001, ****p<0.0001; NS, not significant. Source Data for this Figure could not be made freely accessible, but have been deposited to the controlled-access repository Zenodo.

To experimentally assess the effector potential of ABC subsets, we examined their capacity to differentiate into antibody-secreting cells using a single B cell culture assay. CD11c+ mature B cells from healthy donors and malaria patients were sorted by CD24 and CD27 expression into the four ABC subsets (**Fig. 7D**) and cultured at single-cell density with IL-2, IL-21, and irradiated CD40L-expressing feeder cells for 14–17 days. Supernatants were screened for IgM, IgG, and IgA by ELISA across 1,450 wells (**Fig. 7E**). Of the antibody-positive wells, 37.7% were derived from Early-ABCs (CD24+CD27+), 25.6% from Int-ABC2 (CD24+CD27−), 22% from Int-ABC1 (CD24−CD27+), and 14.7% from Late-ABCs (CD24−CD27−), with a similar distribution in healthy donors and malaria patients (**Fig. S9A-B**). Although all subsets were capable of antibody secretion, the efficiency declined progressively with differentiation stage, consistent with the antibody secretion module scores derived from the transcriptomic data. Surprisingly, many wells produced IgA antibodies, with a small proportion of wells (2-8%) producing more than one antibody isotype (**Suppl. Fig. 9C**). This was likely due to class-switching driven by CD40L and IL-21, as has been described before (*40*).

We next assessed the capacity of ABC subsets to bind and potentially take up microbial particles. Total B cells isolated from PBMCs of seven healthy blood donors were co-cultured with fluorescently labeled *E. coli* bioparticles with interactions assessed by flow cytometry (**Fig. 7F and Fig. S9D-E**). Total CD11c+ B cells showed significantly greater fluorescent particle association than CD11c− B cells. Within the CD11c+ compartment, particle association was highest in Late-ABCs (CD24−CD27−) and lowest in Early-ABCs (CD24+CD27+), consistent with the transcriptomically predicted gradient. Of note, flow cytometry cannot discriminate between true intracellular uptake and surface binding of fluorescent particles; these data should therefore be interpreted as reflecting differential particle association rather than confirmed phagocytosis.

TNFα and IL-10 represent opposing pro-inflammatory and regulatory B-cell effector functions and together provide a concise readout of whether ABCs are functionally polarized towards inflammation or immune regulation. We next assessed TNFα and IL-10 cytokine producing capacity of ABCs by stimulating total B cells isolated from PBMCs of eight healthy blood donors with anti-Ig+CD40L, CpGC (TLR9 ligand), R848 (TLR7/8 ligand), or PAM2CSK4 (TLR2 ligand) for 20 h, followed by 4 h of PMA/ionomycin stimulation in the presence of brefeldin A and monensin. The PMA/ionomycin was added to assess maximum cell function and the impact when primed by different stimuli, as done before (*44*). TNFα and IL-10 production was measured by intracellular cytokine staining (see **Fig. S9F-H** for gating strategies). In all but one condition, CD11c+ B cells produced significantly more cytokines than CD11c– B cells, with TNFα being detected in a larger proportion of cells than IL-10 (**Fig. 7G-H**). Within the CD11c+ compartment, anti-Ig+CD40L drove TNFα production most prominently in CD27-expressing subsets (Early-ABC and Int-ABC1), while CpG induced TNFα production across all subsets. A non-significant decreasing trend in TNFα production was observed across the differentiation axis regardless of stimulus (**Fig. 7I**). IL-10 followed a broadly similar pattern at lower frequencies (**Fig. 7J**).

Together, these findings demonstrate that the ABC differentiation axis is associated with a progressive functional reprogramming: early stages retain plasma cell differentiation potential and cytokine secretion capacity consistent with an activated effector-like state, while late stages acquire particle-associating capacity and antigen presentation features. These data provide functional validation of the transcriptomically inferred module scores and reinforce the biological significance of the CD11c/CD24/CD27 differentiation framework.

## Discussion

Studies investigating ABCs commonly use gating strategies that excludes non-switched or early and intermediate ABC (e.g. CD21–CD27– or IgD–CD27–) and can give the impression of ABCs as a fixed, discrete subset rather than a continuum. In this study, we used an inclusive gating approach based on CD11c to capture a more extended breadth of ABC-like B cells, including potential subsets overlooked by conventional definitions. We selected CD11c because it marked the transcriptionally defined ABC clusters and most proliferating ABCs while remaining largely absent from CD27–CD21+IgD+ naïve B cells, indicating a predominantly memory (CD27+) origin.

We found that ABCs form a heterogeneous continuum shaped by origin, isotype, and tissue environment, spanning early to late stages with gradual transcriptional, phenotypic, and functional shifts across malaria patients and healthy donors. ABCs originated from both IgD–IgM– switched and IgD–IgM+ non-switched GC-derived memory B cells and from IgM+IgD+ marginal zone-like non-switched memory cells of likely extrafollicular origin. CD24 and CD27 resolved intermediate states and linked differentiation stage to antibody-secreting cell capacity, which diminished in later subsets, at least under T cell-dependent stimuli. To comprehensively assess cytokine production across subsets, cells were stimulated with both BCR ligands and TLR agonists, as ABCs have been described as BCR-hyporesponsive (*45*) and proposed to function as innate-like cells (*46*). This revealed a broader functional gradient along the differentiation axis, with early subsets enriched for cytokine secretion capacity and late subsets acquiring antigen presentation and microbial particle association features, establishing stage-dependent functional roles across the ABC continuum. This CD11c/CD24/CD27 organization was conserved between blood and tonsil, and ABC-associated markers were strongly upregulated among proliferating cells, indicating activated rather than quiescent states.

Prior work from us (*16*) and others (*9, 31, 36*) have reported ABC heterogeneity across ages and infections. Sutton et al. investigated B cells in malaria (*9*) and combined CD21 and CD27 with CXCR3 and CD11c. They used pseudotime and clonal analyses of antigen-specific B cells (PfCSP, PfMSP1, and tetanus toxoid specific) and proposed that ABCs branched from GC derived memory. We confirmed similar GC derived ABCs but additionally identified subsets with low or absent somatic hypermutation, many with non-switched BCRs, indicating contributions from non GC origin. Sutton et al. concluded that ABCs represent an alternative lineage rather than simply recently activated B cells (*9*). Our findings support ABCs as both an activated state and a distinct lineage, with maturation stages detectable under homeostatic conditions up to one year after infection, without documented antigenic re-exposure.

Holla et al. (*31*) highlighted that ABC expansion in malaria, HIV, and autoimmunity was strongly influenced by inflammatory and activation signals. Their single-cell and trajectory analyses positioned ABCs as a terminal IFN-γ–enriched state, driven by sustained IFN γ and BCR/TLR cues, that responded to membrane-associated but not soluble antigen. Our data refine this view: rather than a single terminal branch, we resolve internal organisation along a CD24/CD27 axis with two origin-dependent intermediate stages. Consistent with origin-specific signaling, anti-Ig/CD40L drove TNFα most strongly in CD27-expressing subsets while CD27-negative subsets relied more on TLR9 (CpG) responsiveness, suggesting the reported BCR hyporesponsiveness is subset-specific and tied to precursor origin. Together with the inverse gradients of antibody-secretion and particle-association capacity, this indicates a stage-dependent reprogramming from an effector-like early state to a scavenger/antigen-presenting late state.

Other studies specifically investigated the role of ABCs in the immune response against malaria. A recent report by Hopp et al. (*47*) identified PfAMA1 and PfMSP1 specific ABCs as having a CD21–CD27– phenotype, a definition that primarily encompasses late ABC populations. They further showed that the ABCs shared clonal relationships with memory and activated B cells, with the ability to differentiate into antibody-secreting cells in the presence of T follicular helper cells. Similar work by Kochayoo et al. (*48*) defined *P. vivax*–specific ABCs as CD21–CD27– and demonstrated their ability to produce broadly neutralizing antibodies against PvDBPII variants. A separate study by Partey et al. (*49*) examined ABC frequency in malaria-immune adults using a similar gating strategy and found correlations with broader antibody responses, although these antibodies exhibited reduced avidity. These studies establish that ABCs contribute to humoral immunity, however their CD21–CD27–gating captures mainly late subsets. Our data confirm the antibody-secreting potential of late ABCs while showing that early and intermediate subsets have even greater capacity, so conventional late-ABC gating overlooks a substantial part of the antibody-producing compartment. Integrating these populations into future studies could clarify which parts of the ABC continuum contribute to antibody secretion *in vivo* and to infection control.

Studies of B cells in tonsils indicate that local cues can shape ABC phenotypes. Prior *in vitro* work using tonsillar B cells showed that sustained BCR engagement combined with TLR9 and IFN-γ drives naïve, and to a lesser extent memory, B cells toward an ABC-like phenotype (high T-bet, CD86, CD95, FCRL5), whereas GC B cells were comparatively refractory (*35*). This aligns with our observation that CD11c+ B cells are rare within ongoing GC reactions. More recent integrated CITE-seq analysis identified ZEB2 as a core regulator of the CD11c+ ABC program and linked this state to T follicular helper cell support under persistent antigen exposure (*40*). In our dataset, tissue effects were evident for CD86, and tonsillar ABCs were more class-switched overall. Studies of mouse spleens further indicate that CD11c+ T-bet+ ABC-like B cells are enriched outside GCs, favoring the marginal zone and red pulp (*41*), consistent with extrafollicular, or pre-GC class switching and diversification (*50, 51*).

Beyond blood and tonsil, single-cell studies have identified ABCs across spleen (*52*), lymph node (*53*), bone marrow (*54*), kidney (*55*), liver (*52*), synovial tissue (*56, 57*), and cerebrospinal fluid (*54*). In these datasets, ABCs emerge by unsupervised clustering with a conserved transcriptomic signature (CD11c, FCRL4/5, T-bet, ZEB2), while surface marker intensity varies by tissue context. Consistent with this, we observed heterogeneity in migration- and function-associated proteins between blood and tonsils and between ABC subsets, with ABCs more abundant in blood, spleen, and bone marrow than in lymph nodes and tonsils. Together with ABC positioning at the T-B border and marginal zone, these findings reinforce that tissue cues modulate activation-marker expression without altering the underlying program, consistent with our hypothesis that subset organisation is conserved across compartments but excluded from GC structures.

Taken together, our findings address key questions raised by recent reviews (*14, 15*) and provide a unified framework for interpreting ABC heterogeneity. We reconcile prior labels, such as DN-like, CD21low CD27–, and isotype-defined groups by situating them along a single differentiation axis with defined positions and cellular origins.

This study was conducted in the context of malaria, which served as an in vivo model of acute and resolving inflammation, however, the conclusions are not intended to be malaria-specific but rather to reflect general principles of ABC differentiation. Importantly, we did not enrich for or directly identify antigen-specific B cells, and therefore the analyses represent the broader pool of responding B cells rather than malaria-specific populations. As such, the inferred differentiation trajectories and functional properties should be interpreted as characteristics of the overall activated B cell compartment during infection. Additionally, although patients were longitudinally followed, the possibility of antigenic re-exposure cannot be fully excluded beyond clinically documented malaria reinfection, which may influence the persistence or reactivation of ABC subsets. In addition, SHM analyses were not benchmarked against classical memory B cells or germinal center B cells, and therefore conclusions regarding relative mutation levels across ABC subsets should be interpreted within the context of the analyzed populations only.

In summary, ABCs arise predominantly from memory B cells activated outside GCs. Rather than a single linear trajectory, multiple precursor compartments follow parallel paths that converge on a shared late-stage phenotype, accompanied by progressive functional reprogramming: early stages retain plasma-cell and cytokine potential, while late stages acquire particle-association and antigen-presentation features. This stage-dependent shift provides a mechanistic basis for the conflicting functions attributed to ABCs, as reports emphasising antibody production likely captured earlier stages and those highlighting antigen presentation or innate-like activity more terminal ones. A CD11c/CD24/CD27 combination delineates memory origin and differentiation stage and is conserved across blood and tonsil. Residual within-subset heterogeneity in CXCR3, FcRL5, and T-bet likely reflects further modulation by tissue, disease, and activation history that this framework cannot fully resolve, and characterising it across diseases and tissues is a natural next step. By embedding contrasting ABC data within a continuous differentiation axis, this framework offers a practical reference for future work across biological and clinical contexts.

## Material and method

### Ethical Statement

Patients diagnosed with malaria were included in the study at Karolinska University Hospital and patients diagnosed with hantavirus infection at Umeå University Hospital, following providing written informed consent according to ethical permits for malaria (Dnr 2006/893-31/4, 2013/550-32, 2015/2200-32, 2016/1940/32, 2018/2354-32, 2019-03436, 2020-00859, 2020-04147, 2024-00374-02 approved by Ethical Review Board in Stockholm and the Swedish Ethical Review Authority) and for hantavirus (Dnr. 04-113M and 07-162M approved by the Ethical Review Board at Umeå University). Blood samples and tonsils were collected from routine tonsillectomies following informed consent according to ethical permit 2016-53-31, approved by the Ethical Review Board at Umeå University.

### Study cohorts

The malaria cohort has been described previously (*58*). Study participants were >18 years old and diagnosed with malaria at Karolinska University Hospital. After study inclusion, participants were sampled repeatedly over one year following treatment. For this study, we generated several new datasets and repurposed selected datasets. The repurposed data included flow cytometry data following hantavirus infection (day 6-9 after symptom onset) (*19*) with new analysis for **Supplementary Figure 1** and malaria infection data from (*16*) to support CITE-seq data in Figure 2. B cell CITE-seq data was extracted from sequenced PBMCs from (*37*). See **Supplementary Table 1** for details on the malaria study cohorts.

### Sample preparation

Peripheral blood samples were collected into EDTA vacutainer tubes. Peripheral blood mononuclear cells were isolated using density gradient centrifugation with Ficoll-Paque Plus (Cytiva). Briefly, whole blood was diluted with one volume phosphate-buffered saline (PBS) without magnesium or calcium and layered over 15 mL Ficoll-Paque Plus in 50 mL Leucosep tubes (Greiner Bio-One). After centrifugation for at 800 × *g* for 15 min at room temperature (RT) without break, the mononuclear cell layer was removed and washed twice in PBS supplemented with 2% fetal bovine serum (FBS, ThermoFisher Scientific) (spin 300 × *g*, 8 min at RT). After counting using trypan blue, cells were resuspended in FBS supplemented with 10% dimethylsulphoxide (DMSO, Sigma-Aldrich) and placed inside a CoolCell at – 80°C overnight, after which the cells were transferred to liquid nitrogen for long-term storage.

### Culture media

The culture medium used for all experiments was RPMI, supplemented with 10% heat-inactivated FBS, 2 mM L-glutamine, 100 U/mL penicillin, 100 μg/mL streptomycin, 10 mM HEPES, and 0.05 mM 2-mercaptoethanol (all from ThermoFisher Scientific).

### B cell isolation

B cells were isolated from buffy coats from healthy blood donors using the RosetteSep Human B Cell Enrichment Cocktail and 50 mL SepMate tubes (both from Stemcell Technologies). Briefly, 25 μL of the Cell Enrichment Cocktail was added per mL of blood and incubated for 20 min at RT. The sample was then diluted with two volumes of PBS containing 2% FBS. SepMate tubes were preloaded with 15 mL Ficoll–Paque Plus (Cytiva) and centrifuged at 1000 × *g* for 1 minute at RT. The diluted blood sample was layered onto the Ficoll and centrifuged at 1200 × *g* for 10 min with the brake on. The enriched B cell fraction was carefully transferred to new tubes and washed twice with PBS + 2% FBS (300 × *g*, 10 min, brake set to 5, RT). After the final wash, the cells were resuspended in PBS + 2% FBS and counted using Countess II (Invitrogen) following a 1:1 dilution with 0.4% trypan blue. The cells were centrifuged again and resuspended at 5 × 10^6^ cells/mL in freezing medium (FBS + 10% DMSO) and placed inside a CoolCell at –80 °C overnight, after which the cells were transferred to liquid nitrogen for long-term storage. B cell, purity was assessed by flow cytometry and was on average ± SD 89.8 ± 6.2%

### Isolation of Naïve and Memory B cells

Rosette-Sep isolated total B cells were further sorted into naïve and memory B cell fractions using magnetic bead isolation kits from Miltenyi. Briefly, memory B cells were first isolated by positive selection by incubating total B cells with CD27-microbeads for 15 min followed by passing over an LS column placed in a MACS separator. The column was then removed, and the CD27-labelled fraction eluted, counted, washed and resuspended in compete media. The unlabeled fraction containing naïve B cells was further incubated for 5 min with biotin-labeled antibodies to bind non-B cells followed by a 5-minute incubation with anti-biotin microbeads. The cell suspension was then again passed over an LS column placed in the MACS separator. The unlabeled naïve B cell fraction was removed, counted, washed and resuspended in complete media. Memory B cells had an average ± SD purity of 86.7 ± 7.3% and naïve B cells of 83.9 ± 9.2%.

### Cell Trace Violet labeling of sorted B cells

To monitor cell division, naïve and memory B cells were labeled with Cell Trace Violet (CTV) (Thermo Fisher Scientific). Briefly, cells were resuspended in PBS at a density of 2 × 10 cells/mL. CTV was added at a final concentration of 1 μM and the suspension incubated for 7 min at 37°C in a ventilated incubator. The staining was halted by adding three volumes of FBS, followed by dilution with PBS up to 15 mL. The cells were then centrifuged at 300 × *g* for 5 min, washed once in PBS under the same conditions, and washed again in complete media. Finally, the cells were rested in complete media at 37°C with 5% CO for 1 hour before further stimulation or analysis.

### In-vitro B cell stimulation

Previously sorted total B cells were thawed and transferred to a 15 mL Falcon tube. 13 milliliters of complete media was added dropwise while gently mixing, and the cells were rested on ice for 20 min. The cells were then centrifuged at 300 × *g* for 5 min, washed once in complete media under the same conditions, resuspended in 1 mL of complete media, and counted. Purified B cells were stimulated at a density of 1 × 10 cells in a total volume of 200 μL complete media in 96-well round-bottom tissue-culture plates (Techno Plastic Products). Cells were either left unstimulated (complete media) or stimulated with a mix of α-Ig (IgA+IgG+IgM [H+L] AffiniPureF(ab’)2 Fragment goat-anti-human [Jackson ImmunoResearch Laboratories]) at 1.25 μg/mL, CpG-C (Invivogen) at 2.5 μg/mL, and IFNγ (Peprotech) at 25 ng/mL. Plates were incubated for 2-4 days at 37 °C with 5% CO . For intracellular cytokine staining, cells were stimulated for 20 h with the indicated stimulant, soluble CD40L (sCD40L; 1μg/mL; Peprotech), R848 (5μg/mL; Invivogen) and/or PAM2CSK4 (5μg/mL; Invivogen), followed by a further 4 h in the presence of PMA/ionomycin, brefeldin A and monensin (all Invitrogen; each at the manufacturer’s recommended dilution of 1000× or 500×).

### Single B cell culture assay

Thawed PBMCs from healthy donors (n=3) and malaria patients (n=3, 10 days after treatment) were stained at room temperature for 20 minutes with a panel of fluorescent antibodies targeting surface markers used to identify the ABCs and its intermediates and some ABC characteristics. Antibodies were diluted to their optimal concentrations in 50 µL of Brilliant Stain Buffer (BD Biosciences). Following staining, cells were washed twice with FACS buffer (PBS supplemented with 2% FBS) and sorted using a 5-laser Cytek Aurora sorter (Cytek Biosciences). Cells were sorted into 384-well plates at a density of 2 cells/well in 50 µL of RPMI-1640 medium supplemented with 10% FBS, 5×10³ irradiated 3T3 feeder cells expressing CD40L with 50Gy (provided by John Mascola and Nicole Doria-Rose, NIH, Bethesda, MD,USA), IL-2 (10 ng/mL), and IL-21 (50 ng/mL). Plates were incubated at 37 °C with 5% CO for 14-17 days.

After two weeks, IgA, IgG and IgM levels in culture supernatants were quantified by ELISA. Briefly, 384-well plates were coated overnight at 4 °C with goat anti-human IgA, IgG or IgM antibody (1 µg/mL; Jackson ImmunoResearch). To block the remaining protein-binding sites in the coated wells, plates were covered with PBS containing 1% BSA for 1 hour at room temperature and subsequently washed with PBS + 0.05% Tween-20. Supernatants from CD11c+ B cells culture, diluted 1:5 in PBS + 0.1% BSA, were added to the coated plates and incubated for 1.5 hours at room temperature. After washing, HRP-conjugated secondary antibodies (0.04 µg/mL) were added and incubated for 1 hour at room temperature. The following antibodies were used for isotype detection: anti-human IgG-HRP (Jackson ImmunoResearch), anti-human IgA-HRP (Sigma-Aldrich), and anti-human IgG+IgM-HRP (Jackson ImmunoResearch) for IgG, IgA, and IgM, respectively. Plates were washed again, then 20 µL of TMB substrate (ThermoFisher) was added to each well and incubated for 5 minutes at room temperature. The enzymatic reaction was stopped with 50 µL of 1 M sulfuric acid (H SO), and absorbance was immediately measured at 450 nm using a Multiskan SkyHigh spectrophotometer (ThermoFisher).

### Flow cytometry

Cells were washed twice with PBS and incubated on ice for 20 min with Live/Dead stain (ThermoFisher Scientific) diluted 1:1200 in PBS. After staining, cells were washed twice with PBS + 2% FBS before being incubated on ice for 20 min with a mix of antibodies targeting surface markers. For intracellular staining, cells were washed once after surface staining, then incubated at RT for 30 min with FoxP3 Fix/Perm buffer (eBioscience). Following fixation and permeabilization, cells were washed in FoxP3 Wash/Perm buffer and incubated for another 30 min at RT with a mix of antibodies targeting intracellular markers (see **Supplementary Table 2** for antibodies). The cells were then washed again with Wash/Perm buffer, followed by a final wash in PBS + 2% FBS. Finally, samples were resuspended in 300 μL PBS + 2% FBS containing 5 μL CountBright Absolute Counting Beads (ThermoFisher Scientific). For data analysis, samples were processed using either the BD Fortessa or the Cytek Aurora (Cytek Biosciences).

### Targeted single-cell-multi-omics sequencing using the BD Rhapsody platform

We used a previously reported targeted single-cell multi-omics dataset, generated on PBMCs from malaria-infected individuals using the BD Rhapsody platform (*37*). In short, CD45+ PBMCs from 4 individuals (2 primary infected, 2 previously exposed) with three time points (Acute, day 10, 1 year) were profiled on a transcriptional level using the Immune Response Panel (targeting 399 genes), as well as on a cell surface level using 29 AbSeq oligo-conjugated antibodies (BD Biosciences). A total of 73,121 cells were annotated based on the expression of 374 genes and 29 surface markers, resulting in 27 immune cell subsets as previously described (*37*). Here, we digitally isolated all B cells (n=2601 cells) and refined the B cell subsets after re-integration and re-clustering based on RNA data using the Seurat pipeline (v4). RNA counts were normalized using the LogNormalize method with a scale factor of 10,000, and surface protein counts were normalized using centered log-ratio (CLR) normalization. Data were then scaled across all detected genes using the ScaleData function. Formal computational doublet detection was not re-applied to the B cell subset, as the parent dataset had undergone prior quality control as previously described (*37*). Contaminating non-B cell populations were identified by aberrant co-expression of lineage-exclusive markers (CD3E, NKG7, GZMB, PRF1, CD163) and excluded prior to downstream analysis Principal component analysis (PCA) was performed on all scaled genes, and the number of significant principal components was determined by visual inspection of an ElbowPlot of the Harmony-corrected reduction; 10 principal components were retained for initial B cell clustering and UMAP embedding, and 15 principal components for subsequent re-clustering after removal of contaminating populations. Batch correction across donors was performed using Harmony (group.by = sample identity). Unsupervised clustering was performed using the Louvain algorithm at resolution 0.8. Differential gene expression between clusters was assessed using the Wilcoxon rank-sum test implemented in FindAllMarkers and FindMarkers, with a minimum fraction of expressing cells of 25% (min.pct = 0.25) and a minimum log fold-change threshold of 0.25 (logfc.threshold = 0.25). For downstream analyses including Venn diagram comparisons, differentially expressed genes were considered significant at an adjusted p-value ≤ 0.05 and |avg_log FC| > 1. Pseudotime analysis was performed using the Slingshot package (v2.0.0), applying default parameters to clusters derived from Seurat clustering. The CD21 naïve-like B cell cluster was designated as the root, enabling trajectory inference across the identified B cell subsets and capturing the continuum of differentiation pathways.

### 5’V(D)J sequencing using the 10X Genomics platform

PBMCs were thawed from one donor with malaria two weeks after treatment initiation and washed once in complete media and once in PBS before incubation for 20 min on ice with Aqua viability dye (ThermoFisher). The cells were then washed once with FACS buffer (PBS+2% FBS) and resuspended in 100 <u>μ</u>l FACS buffer containing anti-CD3 and anti-CD14 FITC (Dump), anti-CD19 PE-Cy7 and anti-CD11c BV786 (all antibodies from BD Biosciences) and incubated for 20 min on ice. Live, Dump negative, CD19high CD11c+ cells were bulk sorted into PBS supplemented with 0.04% BSA and transferred to the Eukaryotic Single Cell Genomics Facility (ESCG) at SciLifeLab, Stockholm, Sweden. ESCG performed B/T cell enrichment plus cDNA synthesis using the 10X Genomics VDJ Kit. Following quality control, paired-end sequencing (2×50bp, NovaSeq S1) was done for gene expression and (2×150bp, MiSeq) for VDJ analysis. Following sequencing the data was demultiplexed, quality controlled and annotated using the Cell Ranger pipeline version 3.1.0. The gene expression library estimated 1339 cells with 2242 median genes per cell. The VDJ library estimated 1474 cells with 1258 productive V-J spanning pairs.

### Analysis of the 10X genomics gene expression dataset

Single-cell suspensions were processed using the 10X Genomics Chromium platform following the manufacturer’s guidelines. Gene expression libraries were prepared using the 5’ Gene Expression Kit (10X Genomics). The libraries were sequenced on an Illumina NovaSeq S1 system with paired-end reads to achieve a targeted depth of approximately 450,000 reads per cell. Raw sequencing data were processed using Cell Ranger (v3.1.0) (10X Genomics) for demultiplexing, alignment to the GRCh38 reference genome, and generation of feature-barcode matrices. The resulting gene expression data were imported into Seurat (v5.2.1) for downstream analysis. Quality control was performed by retaining cells with between 500 and 3,500 detected genes, mitochondrial gene expression below 20%, and ribosomal gene content above 10%. The upper threshold on detected genes (< 3,500) additionally served as a conservative proxy for doublet exclusion, as cell multiplets characteristically exhibit abnormally elevated gene counts relative to singlets. Formal computational doublet detection was therefore not applied separately; furthermore, given that this dataset represents an enriched B cell population with low expected cellular heterogeneity, residual homo-doublets would not be expected to generate artifactual transcriptional clusters. Prior to scaling, hemoglobin genes (HBB, HBA1, HBA2, HBG1, HBG2, HBD), the long non-coding RNA MALAT1, all mitochondrial-encoded transcripts (MT-*), and ribosomal protein-coding genes (RPL/RPS family) were removed to prevent their disproportionate influence on dimensionality reduction and clustering. Data normalization was performed using the LogNormalize method (scale factor = 10,000). The top 2,000 highly variable genes were identified using the variance-stabilizing transformation (VST) method and used as input for principal component analysis (PCA). The number of statistically significant principal components was determined by the JackStraw procedure (100 replicates), corroborated by visual inspection of an ElbowPlot; 20 principal components were retained and applied consistently for nearest-neighbor graph construction (FindNeighbors), unsupervised clustering (FindClusters, resolution = 0.8), and UMAP embedding. Differential gene expression between clusters was assessed using the Wilcoxon rank-sum test (FindAllMarkers), with a minimum fraction of expressing cells of 25% (min.pct = 0.25) and a minimum log fold-change threshold of 0.25 (logfc.threshold = 0.25). Per-cell immune function scoring was performed using the UCell package with curated gene signatures covering cytokine production, phagocytosis, antibody secretion, and antigen presentation; pairwise comparisons between clusters were assessed using Wilcoxon tests with Benjamini-Hochberg correction. Gene set enrichment analysis was performed using fgsea against MSigDB Hallmark gene sets, ranked by cluster-level average log fold-change. Pathway enrichment analysis was performed using the ReactomeGSA package (v1.18.0) in R, applying gene set variation analysis (GSVA) across clusters. Selected pathways related to B cell biology, antigen presentation, complement activation, cell cycle, and cellular activation were visualized using customized GSVA heatmaps. Differentially expressed genes between clusters (adjusted p-value < 0.05, |log2FC| > [X]) identified with Seurat’s FindMarkers function were used as input for pathway analysis using QIAGEN Ingenuity Pathway Analysis (IPA, QIAGEN Inc.). Core analyses were performed for each comparison to identify enriched canonical pathways, upstream regulators, and biological functions, using default settings (Human reference set, [Direct + Indirect / Direct only] relationships, [Experimentally Observed / Experimentally Observed + High confidence predicted] confidence filter). Comparison analysis was then performed across clusters/conditions to identify shared and divergent pathway activity, restricted to immune-related functional categories.

### Fluorescent particle association assay

The capacity of B cell subsets to associate with microbial particles was assessed using the Phagocytosis Assay Kit (Green *E. coli*, ab235900; Abcam/BioVision), which provides heat-killed, fluorescently pre-labeled *E. coli* particles. B cells isolated from PBMCs of seven healthy donors were co-incubated with *E. coli* particles for 1 hou according to the manufacturer’s instructions. Following incubation, cells were washed and surface-stained as described above for flow cytometric acquisition. The percentage of green fluorescent particle-positive cells was quantified within CD11c− and CD11c+ B cell fractions and further resolved by CD24 and CD27 expression. As no cytochalasin D inhibitor control was included to block active internalization, the assay measures total fluorescent particle association (surface binding and/or internalization) rather than confirmed phagocytosis, and results are interpreted accordingly.

### Analysis of the 10X Genomics 5’V(D)J library dataset

B cell clonal analysis was performed using the Immcantation framework and enclone. V(D)J gene annotation was done with IgBLAST and processed with Change-O. Clonal lineages were assigned by grouping sequences with the same V and J genes, the same CDR3 length, and at least 82% sequence similarity in the CDR3 region, using a distance threshold of 0.18 estimated by findThreshold() from scoper. Diversity analysis, V(D)J gene usage, and CDR3 length distributions were analyzed with alakazam. Clonal lineage trees were visualized using dowser. To visualize clonal expansion at the single-cell level, we used enclone to create honeycomb plots (honeymaps), showing clonal distributions and isotype switching across the 10X Genomics data.

### Statistical analysis

Flow cytometry data were analyzed using GraphPad Prism version 10.4.2. Sequencing data was analyzed using R-studio. Group comparisons were performed using One or Two-way ANOVA with Geisser-Greenhouse correction followed by Sidak’s or Tukey’s posttests. Significance was defined as ns=p>0.05, *p<0.05, **p<0.01, ***p<0.001, ****p<0.0001.

## Code and data availability

The R code describing the single-cell RNA sequencing analysis is available via GitHub (https://github.com/SundlingLab/ABCdifferentiation). European Data Regulations preclude open deposition of sensitive personal data into public repositories. This includes biological data that can be traced back to an individual, so-called pseudoanonymized data. We have therefore made the transcriptomic and other source data available pending relevant permits via Zenodo (doi:10.5281/zenodo.20814279). This includes a metadata record describing the existing datasets.

## Acknowledgements

We would like to thank all study participants for contributing to the study and clinicians, nurses, laboratory staff, and students for assisting with study inclusion, sampling, and processing. We thank the Center for Molecular Medicine flow cytometry core and Science for Life Laboratory, the Knut and Alice Wallenberg Foundation, the National Genomics Infrastructure, and Uppsala Multidisciplinary Center for Advanced Computational Science for assistance with massively parallel sequencing and access to the UPPMAX computational infrastructure. The Graphical abstract was made using BioRender.

The study was supported by grants to CS from the Swedish Research Council (2019-0194 and 2023-01943), Åke Wiberg foundation (M18-0076), Magnus Bergvall foundation (2017-02043 and 2018-02656) and Karolinska Institutet (FS-2020:0007 and FS-2022:0010). ADCG and MJL were supported by PhD grants from Karolinska Institutet to CS (2020–00878) and to AF (2019-00992), respectively. The malaria cohort was supported by grants to AF from the Swedish Research Council (521-2012-3311, 2015-02977, 2018-02688, 2018-04468, 2021-03105), Region Stockholm ALF Medicine (20150135, 20180409, 960104, 986923) and the Marianne and Marcus Wallenberg Foundation.

## Author contributions

ADCG and CS conceptualized the study. ADCG, CS, LK, and MP designed and performed experiments. ADCG, CS, LK, MJL, MP, GEP, and ZM analyzed data. ADCG, ZM, MJL, MP and CS generated figures and visualizations. AF and MF provided scientific input and critical reagents. CS provided funding and student supervision. ADCG wrote the initial draft with input from CS and LK. All authors contributed to the revision of the manuscript.

