## Supplemental material for "CD24 and CD27 resolve ontogeny and gradual differentiation of CD11c+ atypical memory B cells after malaria infection"

**Supplementary Table 1. Descriptive statistics of the study participants.**

| **All donors** | **Malaria** | **Healthy** |
|---|---|---|
| Number of participants | 60 | 14 |
| Female sex, n (%) | 16 (42.7) | 10 (71) |
| Age, years, median (range) | 38 (20-78) | 32 (21-57) |
| Parasitemia, % infected RBCs, median (range)[2] | 0.7 (0.009-17.0) | – |
| *Plasmodium falciparum* infection, n (%) | 57 (95) | – |
| Previous malaria exposure, n (%)* | 36 (65) | 0 (0) |
| ***Related to Figure 1 (flow cytometry)*** | | |
| Number of participants | 4 | – |
| Female sex, n (%) | 3 (75) | – |
| Age, years, median (range) | 47 (36-51) | – |
| Parasitemia, % infected RBCs, median (range)[2] | 1 (0.9-1.4) | – |
| *Plasmodium falciparum* infection, n (%) | 4 (100) | – |
| Previous malaria exposure, n (%) | 3 (75) | – |
| ***Related to Figure 2 (flow cytometry)*** | | |
| Number of participants | 18 | – |
| Female sex, n (%) | 4 (22) | – |
| Age, years, median (range) | 34 (27-54) | – |
| Parasitemia, % infected RBCs, median (range)[2] | 0.4 (0.09-8) | – |
| *Plasmodium falciparum* infection, n (%) | 18 (100) | – |
| Previous malaria exposure, n (%) | 13 (72) | – |
| ***Related to Figure 2 (BD Rhapsody)*** | | |
| Number of participants | 4 | – |
| Female sex, n (%) | 0 (0) | – |
| Age, years, median (range) | 42 (26-62) | – |
| Parasitemia, % infected RBCs, median (range)[2] | 0.7 (0.1-1.4) | – |
| *Plasmodium falciparum* infection, n (%) | 4 (100) | – |
| Previous malaria exposure, n (%) | 2 (50) | – |
| ***Related to Figure 3/4/5 (flow cytometry + 10X)*** | | |
| Number of participants | 9 | 6 |
| Female sex, n (%) | 5 (56) | 5 (83.3) |
| Age, years, median (range) | 46 (22-78) | 32 (24-46) |
| Parasitemia, % infected RBCs, median (range)[2] | 1.2 (0.1-7.0) | – |
| *Plasmodium falciparum* infection, n (%) | 6 (67) | – |
| Previous malaria exposure, n (%)* | – | 0 (0) |
| ***Related to Supplementary Figure 4*** | | |
| Number of participants | 51 | 14 |

| | | |
|---|---|---|
| Female sex, n (%) | 11 (22) | 10 (71) |
| Age, years, median (range) | 37 (20-68) | 32 (21-57) |
| Parasitemia, % infected RBCs, median (range)[2] | 0.6 (0.009-17) | – |
| *Plasmodium falciparum* infection, n (%) | 51 (100) | – |
| Origin in malaria endemic area, n (%) | 33 (65) | 0 (0) |

*Missing information from 5 donors

**Supplementary Table 2. Antibody list for flow cytometry and cell sorting panels**

| Antibody specific for | Clone | Fluorochromes | Company |
|---|---|---|---|
| CD1c | F10/21A3 | BB515 | BD |
| CD3 | UCHT1 | APC-R700 | BD |
| CD4 | SK3 | BV750 | BD |
| CD8 | SK1 | PerCp-Cy5.5 | BD |
| CD10 | HI10a | APC-R700 | BD |
| CD11c | B-ly6 | BB515, BV650 | BD |
| CD19 | HIB19 | PE-Cy7, APC-R700 | BD |
| CD19 | SJ25C1 | BUV737 | BD |
| CD20 | 2H7 | APC-H7, BV510 | BD |
| CD24 | ML5 | PE-CF594, PE | BD |
| CD21 | HB5 | PE-Cy7 | ThermoFisher Scientific |
| CD21 | B-ly4 | BV421 | BD |
| CD27 | M-T271 | BV650, BV786 | BD |
| CD38 | HIT2 | PerCp-Cy5.5 | BD |
| CD38 | HIT2 | APC-Fire750, APC-Fire810 | Biolegend |
| CD55 (DAF) | IA10 | BB515 | BD |
| CD56 | NCAM16.2 | APC-R700 | BD |
| CD62L | DREG-56 | BV480 | BD |
| CD80 | B7-1 | PE-Cy5 | Thermo Fisher Scientific |
| CD86 | IT2.2 | PE-Cy5 | Biolegend |
| CD85j | GHI/75 | Biotin | BD |
| CD268/BAFF-R | 11C1 | Alexa Fluor 647 | BD |
| CCR1 | 53504R | PE-Cy5.5 | Novus |
| CXCR3 | 1C6 | PE-Cy5, PE-Cy7 | BD |
| FcRL3 | H5/FcRL3 | Alexa Fluor 647 | Biolegend |
| FcRL4 | A1 | BUV661 | BD |
| FcRL5 | 509f6 | PE/RB613 | Biolegend |
| Ki67 | B56 | BV421 | BD |
| IgD | IA6-2 | BB515, BB700, FITC | BD |
| IgM | G20-127 | BV605 | BD |
| IL10 | JES3-9D7 | R718 | BD |
| NKG7 | 2G9A10F5 | PE | Beckman Coulter |
| T-bet | O4-46 | Alexa Fluor 647 | BD |
| TNF-alpha | MAb11 | BV650 | BD |
| Cell trace violet | | | Thermo Fisher Scientific |
| Live/dead | | Blue, Green | Thermo Fisher Scientific |
| Streptavidin | | BUV395 | BD |

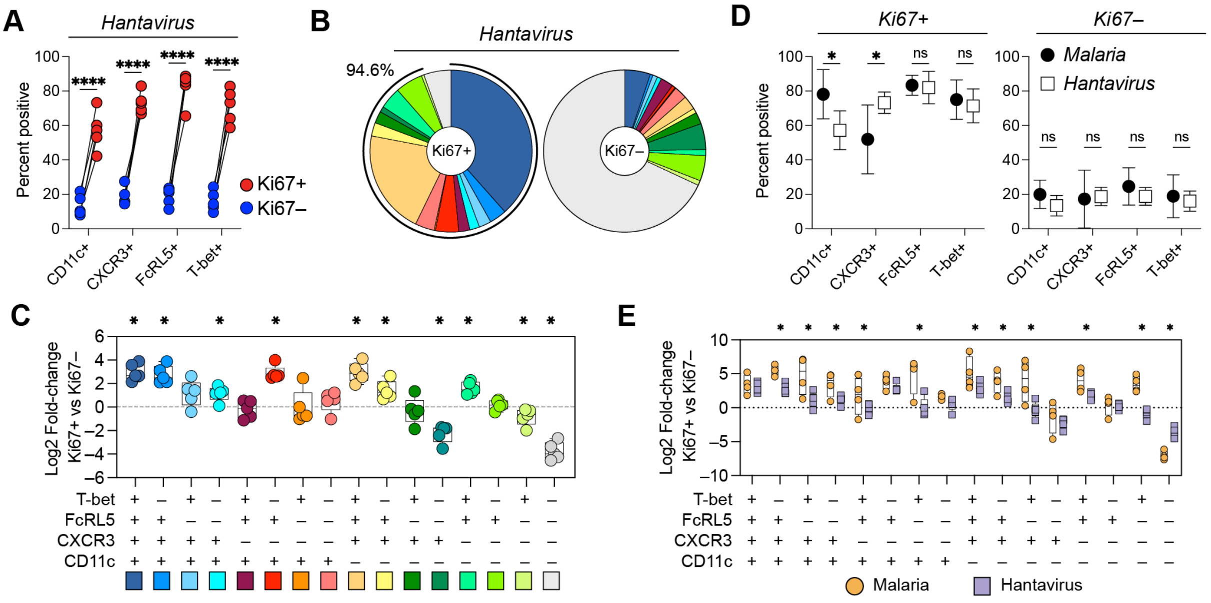

***Supplementary Figure 1. Diversity of atypical B cell-associated markers on B cells after Hantavirus infection**. (**A**) Frequency of mature B cells positive for CD11c, T-bet, CXCR3, and FcRL5 among Ki67+ and Ki67– cells in patients with the hantavirus PUUV. Statistics was evaluated using a paired two-way ANOV followed by Sidak's post-hoc test, ****p<0.0001. (**B**) Donut plot showing the frequencies of populations expressing one or several ABC-associated markers among Ki67+ (left) or Ki67– (right) cells in Hantavirus infection (n=5). (**C**) Log2 fold change representing the expansion of each population between Ki67+ and Ki67– cells in individuals with Hantavirus infection. Statistics evaluated using ratio paired t-tests between KI67+ and KI67– frequencies for each marker combination prior to fold-change calculation. *p<0.05. (**E**) $Log_2$ fold-change ($Ki67^+$ vs $Ki67^-$) for each marker combination, compared between malaria (orange circles) and hantavirus (purple boxes). Differences were tested by two-way ANOVA with FDR correction; *p<0.05.*

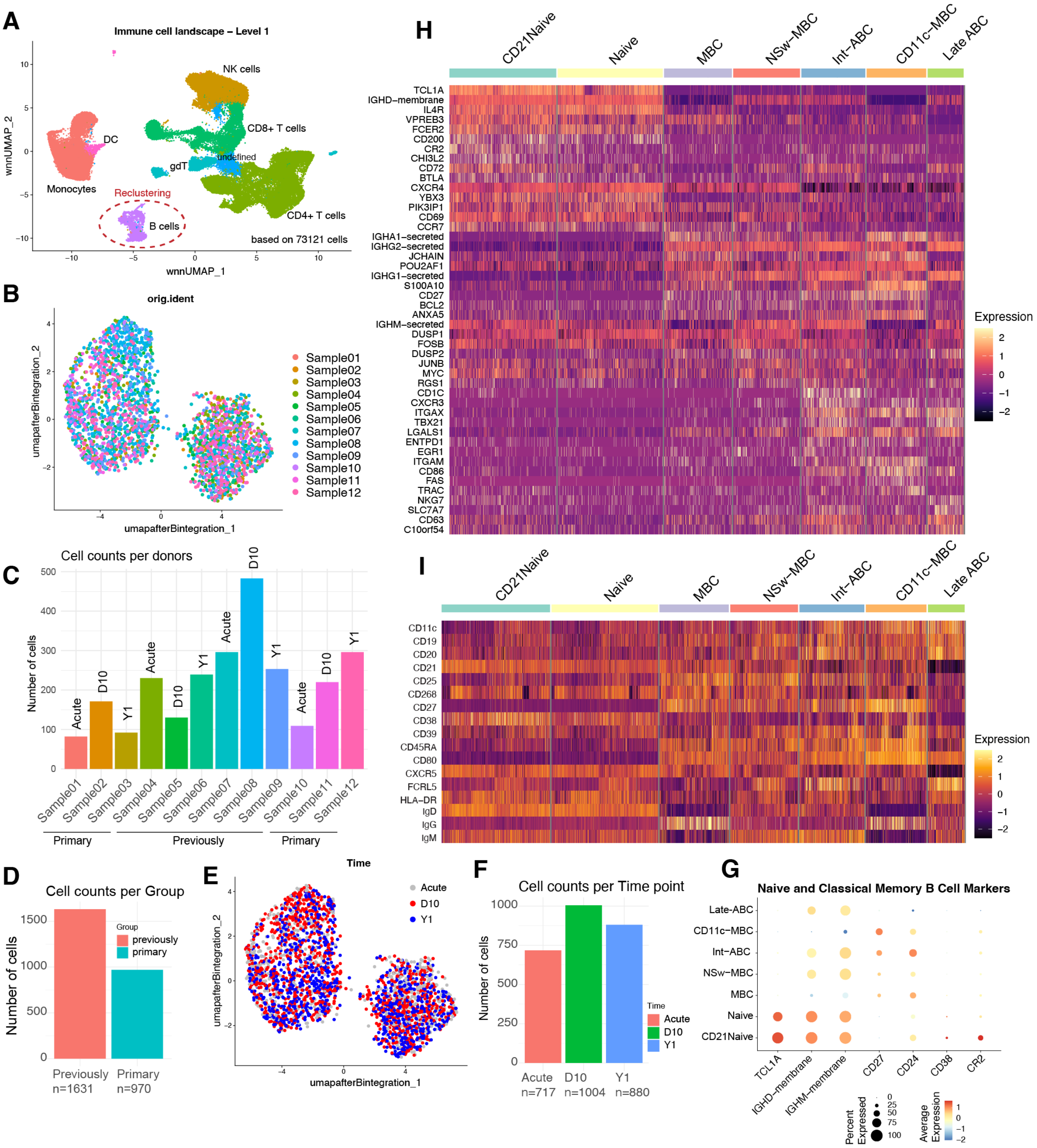

***Supplementary Figure 2. Longitudinal single-cell transcriptomic and CITE-seq analysis after acute malaria. (A)** UMAP clustering of all immune cells from longitudinal samples collected from malaria-infected patients. **(B)** UMAP re-clustering of the B cell compartment, colored by patient of origin. **(C)** The number of cells for each donor and time-point. **(D)** The number of cells for previously exposed and primary infected individuals. **(E)** UMAP re-clustering of the B cell compartment, colored by sampling time point (acute, 10 days post-infection, and 1-year post-infection). **(F)** The number of cells at each time-point. **(G)** Dot plot showing the expression of naive and memory B cell markers across all B cell clusters. **(H)** Heatmap of the top differentially upregulated genes across B cell clusters. **(I)** Heatmap of surface protein marker expression across all B cell clusters.*

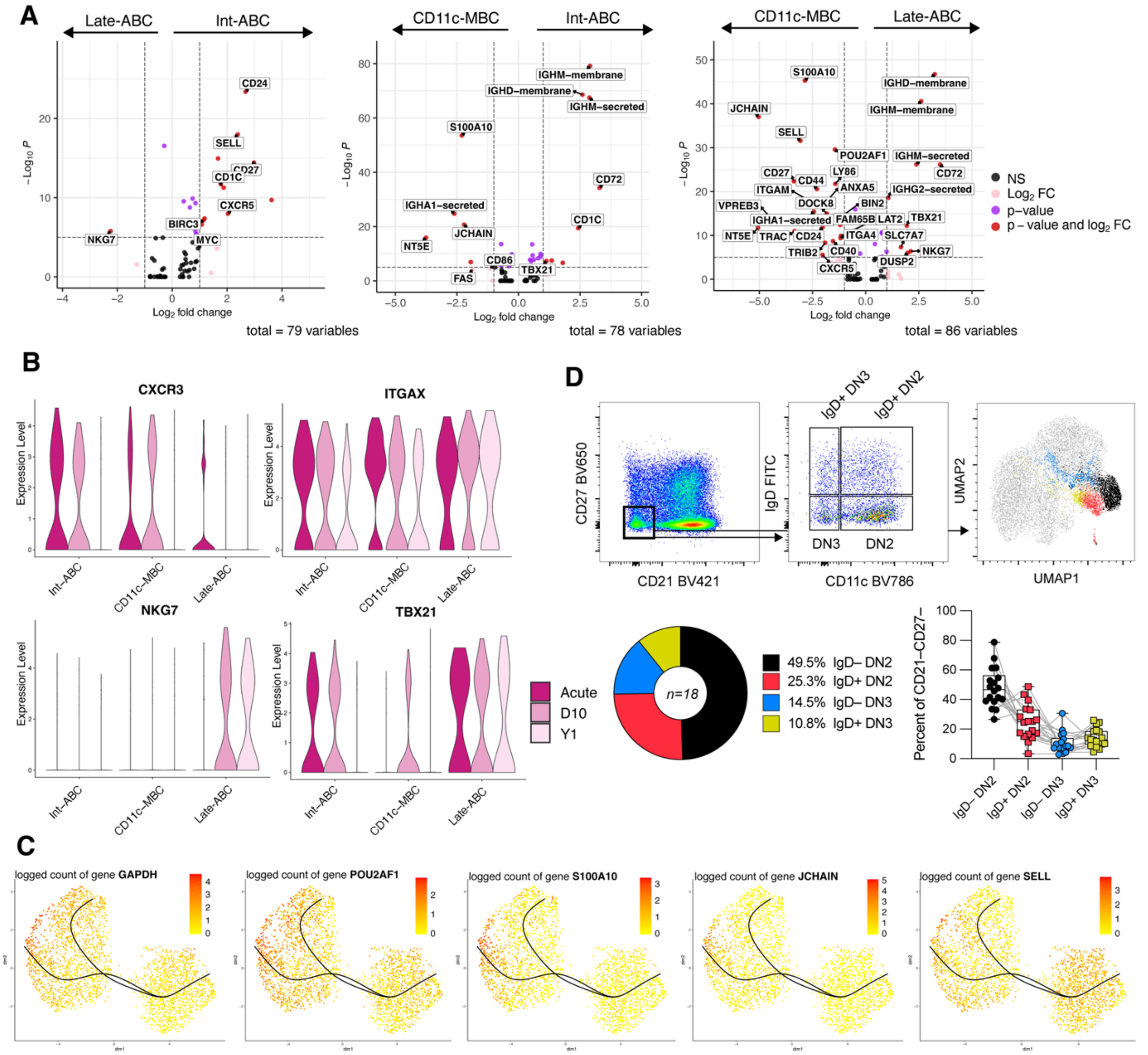

***Supplementary Figure 3. Differential expression and pseudotime analysis of longitudinal single-cell transcriptomic and CITE-seq analysis after acute malaria.***

***(A)*** *Volcano plot comparing the three atypical-like clusters (CD11c+ MBC, Int-ABC, and Late-ABC), highlighting differentially expressed genes.* ***(B)*** *Violin plot showing the expression of atypical B cell markers over time in the atypical-like clusters.* ***(C)*** *UMAP colored by Slingshot pseudotime, with smoothed curves representing two inferred lineages and the top five differentially expressed genes at each lineage endpoint. (****D****) Top panels indicate flow cytometry of 63,000 mature B cells concatenated from 18 donors with acute malaria. Gating for CD21– CD27– Late-ABC followed by gating for IgD versus CD11c to identify DN2 and DN3 cells (+/– IgD expression) with populations overlayed in different colors on the full UMAP embedding. Bottom panels indicate the overall (pie-graph) and individual donor (box-plots) frequency of each DN population among Late-ABCs.*

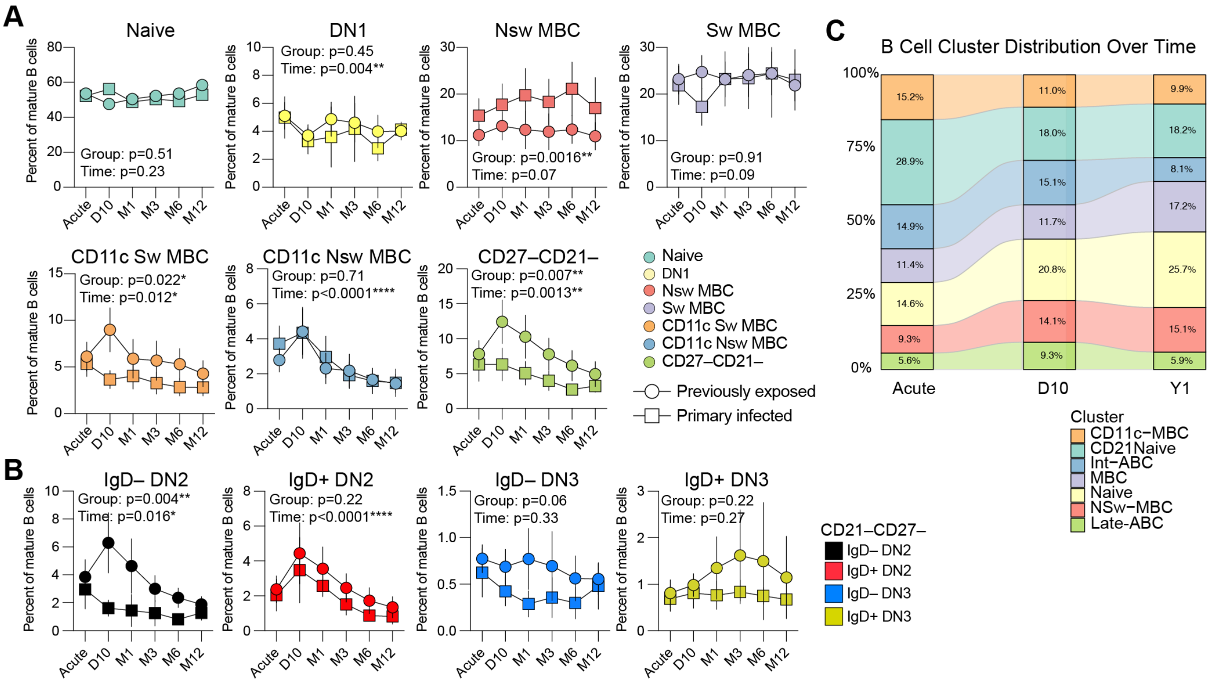

***Supplementary Figure 4. Temporal kinetics of B cell populations up to one year after acute malaria.***

*(**A**) The frequency of Naïve (IgD+CD21+CD27–), DN1 (IgD–CD27–CD21+), Nsw MBC (IgD+CD27+), Sw MBC (IgD–CD27+), CD11c Sw MBC (IgD–CD27+CD11c+), CD11c Nsw MBC (IgD+CD27+CD11c+), and CD27–CD21– B cell subsets was assessed a the time of acute malaria (n=49), and 10 days (n=32), 1 month (n=23), 3 months (n=26), 6 months (n=23), and 12 months (n=27) after treatment initiation. Kinetics for primary infected (boxes) and previously exposed (circles) donors is shown. (**B**) Same analysis as in (A) but for IgD+ and IgD– DN2 (CD27–CD21–CD11c) and DN3 (CD27–CD21–CD11c–) subsets. (**C**) Alluvial plot indicating the frequency of cells within each cluster based on CITE-seq data during Acute malaria, and 10 days, and 1 year after treatment (n=4). Statistical analysis for (A) and (B) was done using a mixed-effects model results for comparisons between primary infected and previously exposed indicated by the Group variable and changes over time indicated by the Time variable. P-values are indicated in the graphs with *p<0.05, **p<0.01, ***p<0.001, ****p<0.0001.*

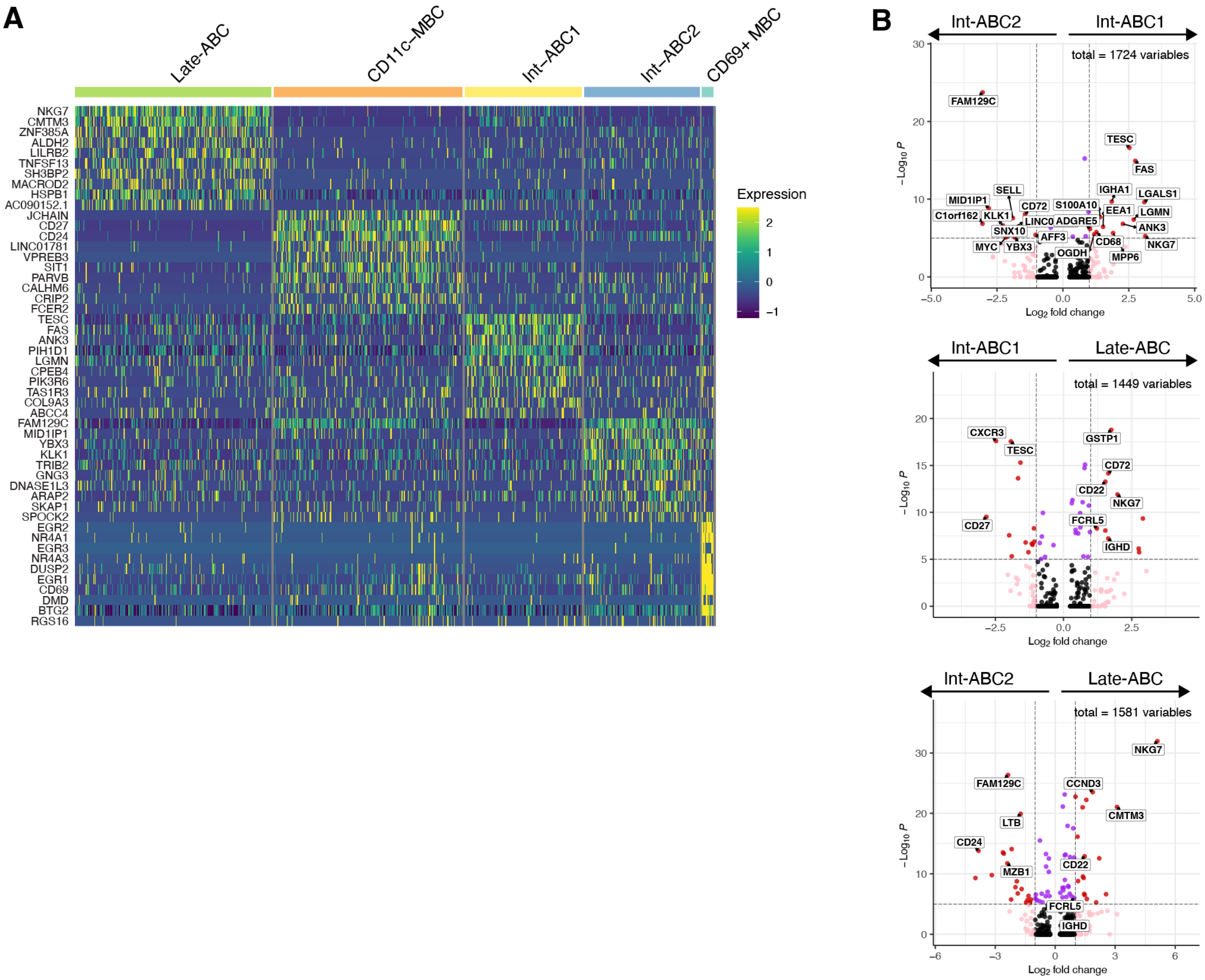

***Supplementary Figure 5. Differential expression analysis of 10X genomics generated single cell transcriptomic data at two weeks after acute malaria.***

***(A)*** *Heatmap of the top 10 differentially expressed genes across all CD11c+ B cell clusters.* ***(B)*** *Volcano plot comparing the three atypical-like clusters (Int-ABC1, Int-ABC2, and Late-ABC), highlighting differentially expressed genes.*

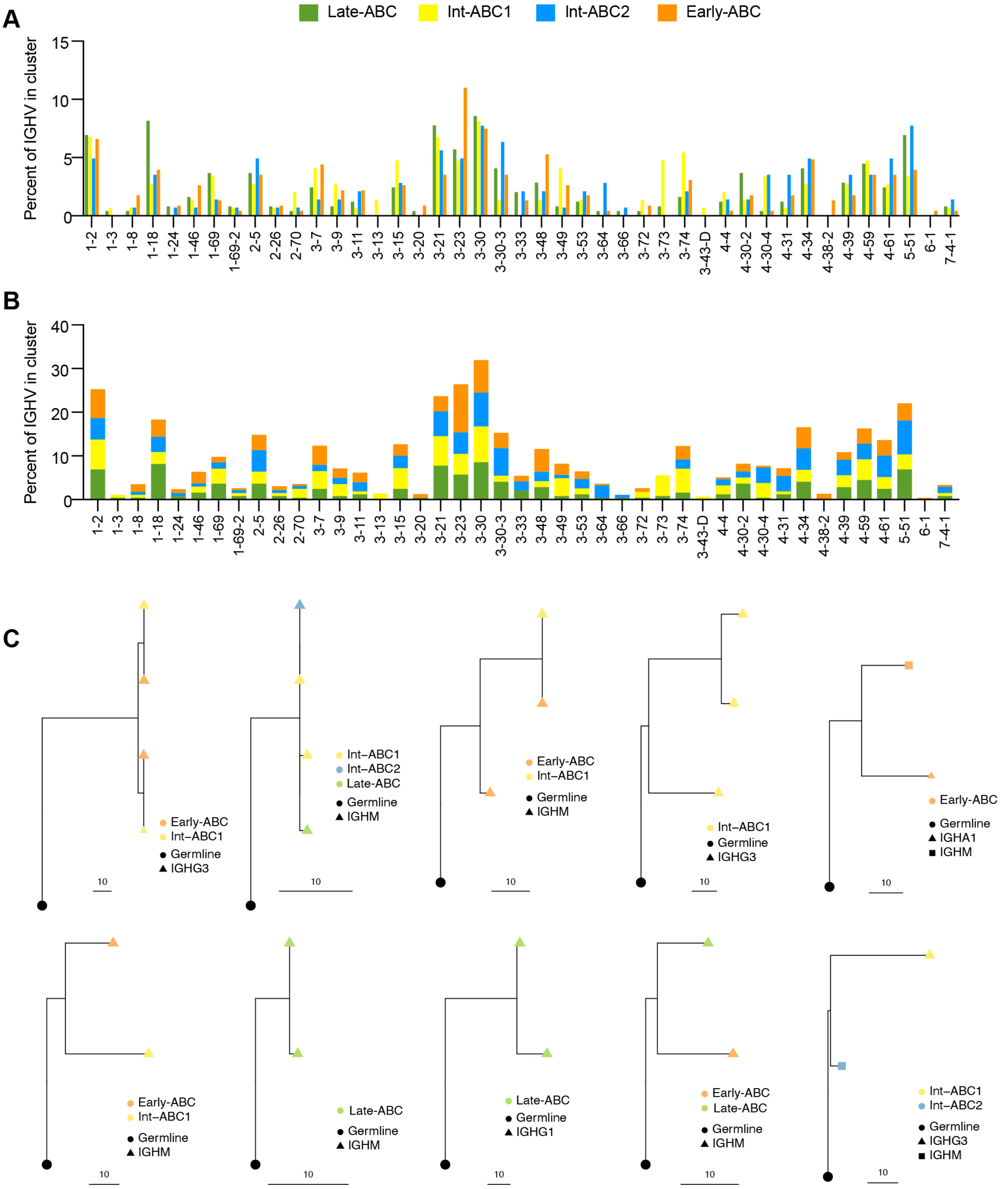

***Supplementary Figure 6. V-gene usage in CD11c+ B cell clusters***.

***(A-B)*** *The frequency of gene usage for each V-gene segment out of all sequenes for each cluster with Late-ABC (green, n=245), Int-ABC1 (yellow, n=147), Int-ABC2 (blue, n=142), and Early-ABC (orange, n=227). The frequencies are shown side by side (****A****) or summarized (****B****). (****C****) Clonal lineage trees based on single cell V(D)J sequence data. Inferred germlines are indicate by black dot. Color indicates which cluster the cells come from. The symbol indicates B cell receptor isotype with the key in the individual lineage trees. Distance is indicated by line corresponding to mutations compared with germline.*

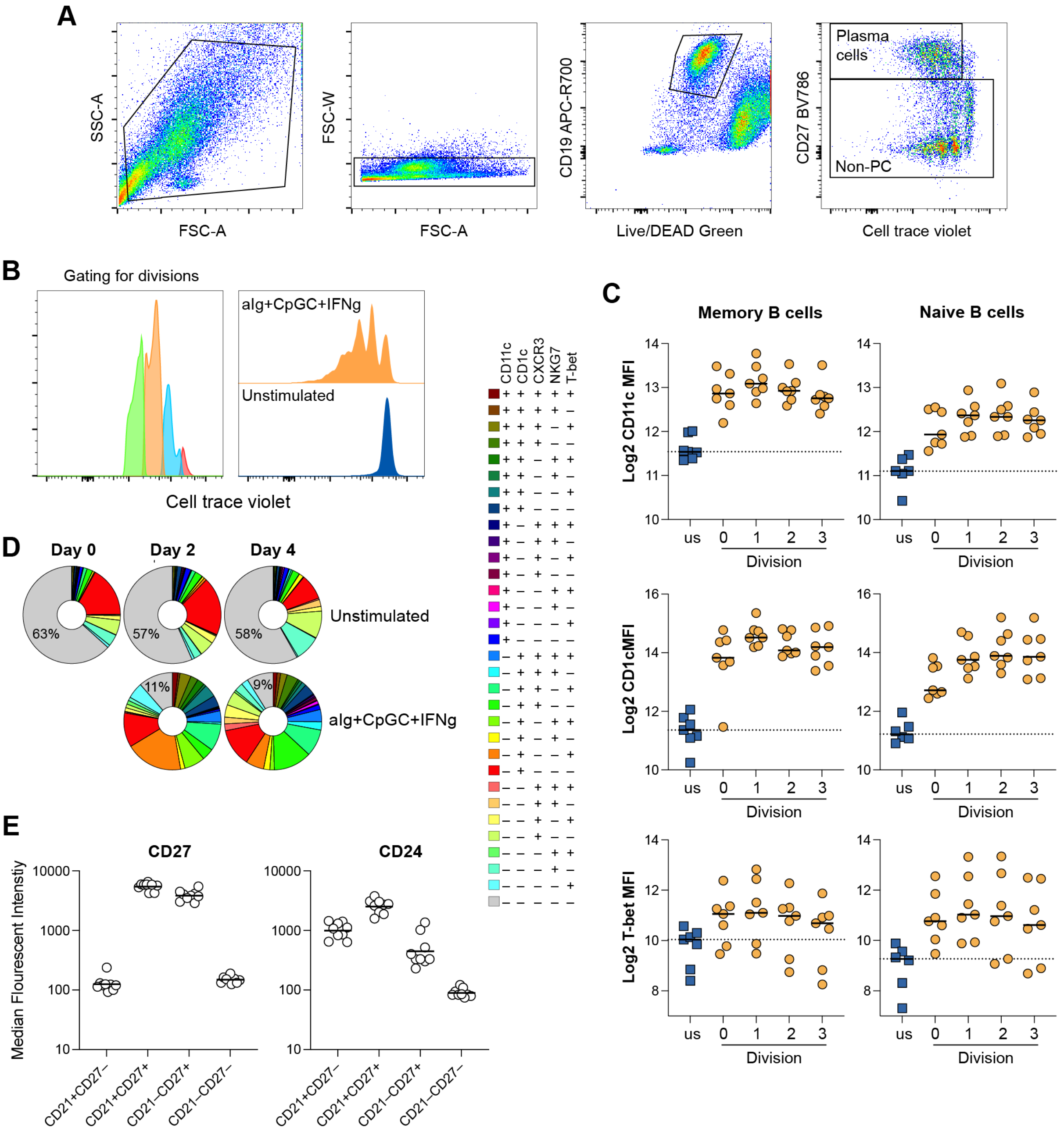

***Supplementary Figure 7. B cell expression after in vitro stimulation or ex vivo after acute malaria.***

*(**A**) Gating strategy to analyze marker expression on non-plasma cells (PC). Example shows sorted memory cells stimualted with aIg+CpGC+IFNg for 4 days. (**B**) Gating of cells after different divisions (left) and example of cell trace violet dilution for stimulated (orange) and unstimulated (blue) cells (right). (**C**) CTV labelled sorted memory and naïve B cells left unstimulated (blue boxes) or stimulated with anti-Ig + CpG-C + IFNg (orange circles) for 4 days. CD11c, CD1c, T-bet median flourescent intensity in undivided and dividing cells with plasma cells gated away. n=6-7 donors pooled from four separate experiments. (**D**) Boolean gating of ABC markers at Day 0, 2, and 4 of stimulation. (**E**) CD27 and CD24 median flourescent intensity in populations gated for CD21 and CD27 from malaria patients (n=9 donors) sampled at the acute time point or 10 days after treatment initiation.*

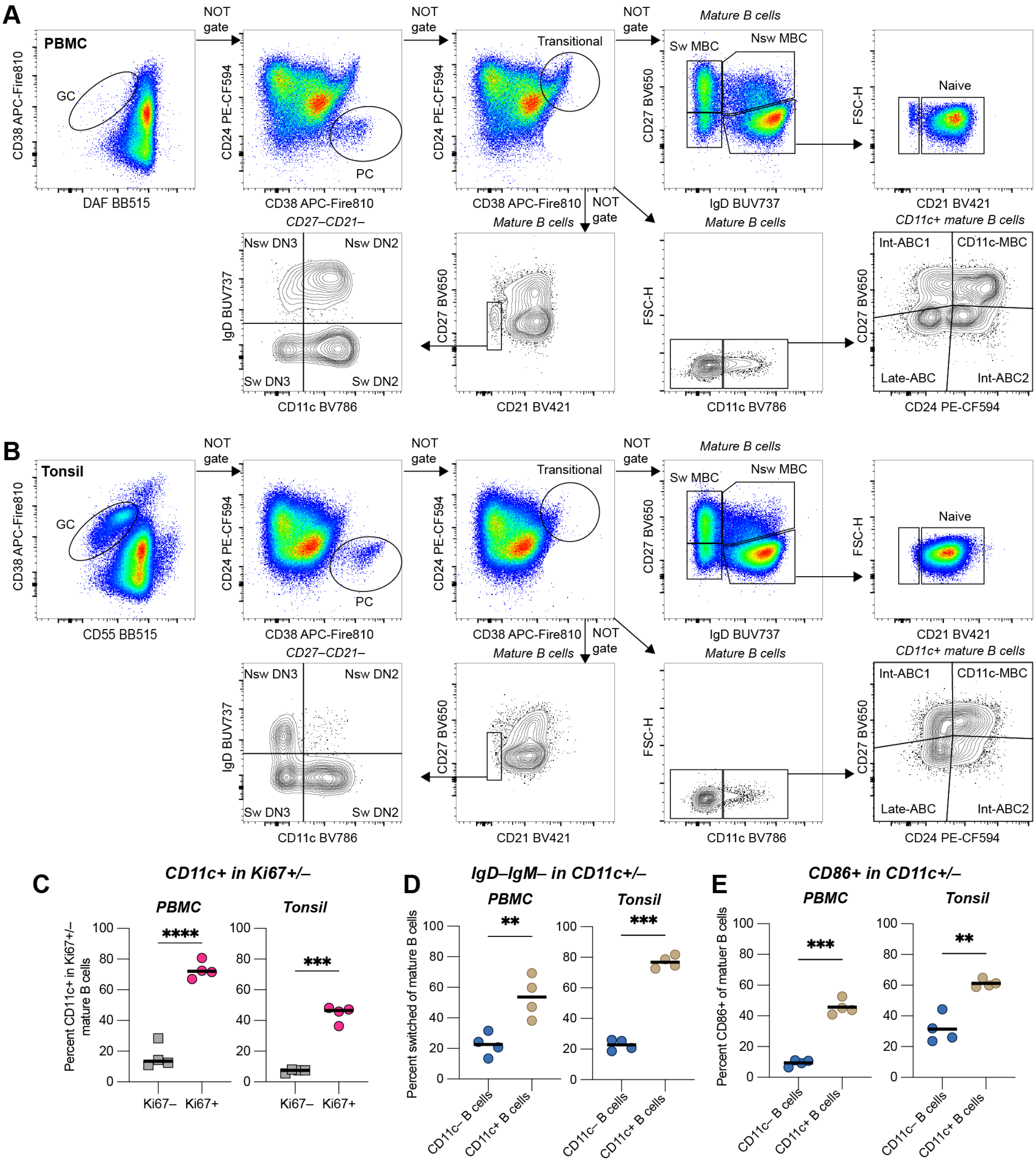

***Supplementary Figure 8. Comparison of B cell populations in matched tonsil and PBMC samples.***

*(**A-B**) Gating strategy for cell populations in PBMC (**A**) and Tonsil (**B**) samples indicated in main Figure 6. FACS plots represent concatenated data from all 4 donors. (**C-E**) Each symbol indicates a donor. Statistical analysis was done by paired t-test with **p<0.01, ***p<0.001, ****p<0.0001. (**C**) Frequency of CD11c+ B cells among Ki67– and Ki67+ mature B cells in PBMC (left, grey box) and Tonsil (right, pink circle) samples. (**D**) Frequency of switched (IgD–IgM–) B cells among CD11c– and CD11c+ mature B cells in PBMC (left, blue circle) and Tonsil (right, yellow circle) samples. (**E**) Frequency of CD86+ B cells among CD11c– and CD11c+ mature B cells in PBMC (left, blue circle) and Tonsil (right, yellow circle) samples.*

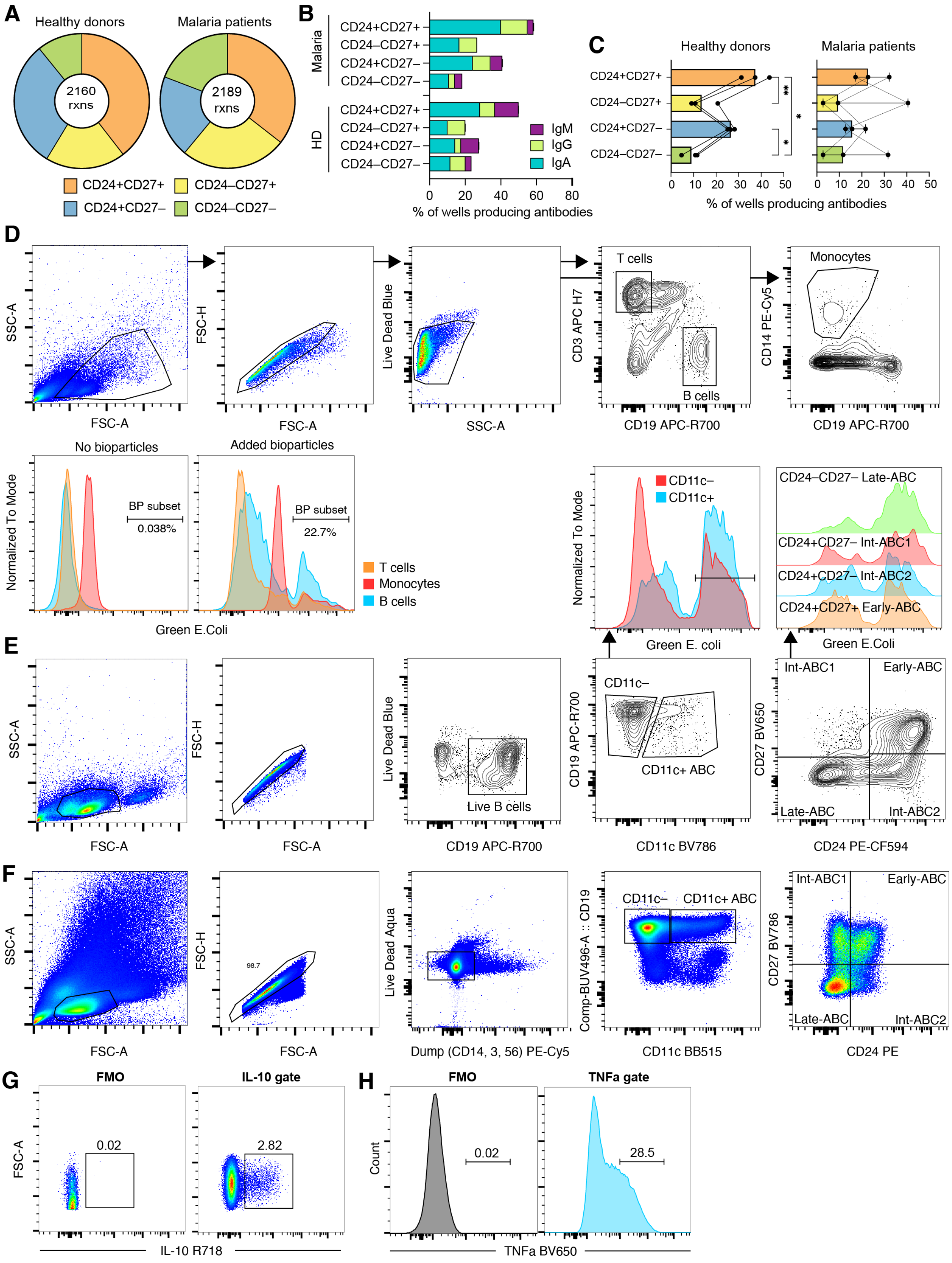

***Supplementary Figure 9. Functional characterization of ABC subsets.***

*(**A**) Overall distribution of antibody positive wells for healthy donors (left) and malaria patients (right). (**B**) Antibody isotype contribution in the different cultured ABC populations, shown for healthy donors and malaria patients. Statistical evaluation by 2-way ANOVA followed by Tukey's posttest. *p<0.05, **p<0.01. (**C**) Comparison between ABC populations for healthy donors (left) and malaria patients (right); statistics as in (B).*

*(**D**) Gating strategy for total T cells, B cells, and monocytes and for bioparticle positive cells. (**E**) Gating strategy for assessing Bioparticle binding in ABCs and ABC subsets. (**F**) Gating strategy for assessing intracellular cytokine production in ABCs and ABC subsets. (**G**) Fluorescence minus one (FMO) to set the gate for IL-10 and (**H**) TNF-alpha.*
